# The influence of inter-regional delays in generating large-scale brain networks of phase synchronization

**DOI:** 10.1101/2023.03.27.534336

**Authors:** N. Williams, A. Ojanperä, F. Siebenhühner, B. Toselli, S. Palva, G. Arnulfo, S. Kaski, J.M. Palva

## Abstract

Large-scale networks of phase synchronization are considered to regulate the communication between brain regions fundamental to cognitive function, but the mapping to their structural substrates, *i.e.*, the structure-function relationship, remains poorly understood. Biophysical Network Models (BNMs) have demonstrated the influences of local oscillatory activity and inter-regional anatomical connections in generating alpha-band (8–12 Hz) networks of phase synchronization observed with Electroencephalography (EEG) and Magnetoencephalography (MEG). Yet, the influence of inter-regional conduction delays remains unknown. In this study, we compared a BNM with standard “distance-dependent delays”, which assumes constant conduction velocity, to BNMs with delays specified by two alternative methods accounting for spatially varying conduction velocities, “isochronous delays” and “mixed delays”. We followed the Approximate Bayesian Computation (ABC) workflow, i) specifying neurophysiologically informed prior distributions of BNM parameters, ii) verifying the suitability of the prior distributions with Prior Predictive Checks, iii) fitting each of the three BNMs to alpha-band MEG resting-state data (*N* = 75) with Bayesian Optimisation for Likelihood-Free Inference (BOLFI), and iv) choosing between the fitted BNMs with ABC model comparison on a separate MEG dataset (*N* = 30). Prior Predictive Checks revealed the range of dynamics generated by each of the BNMs to encompass those seen in the MEG data, suggesting the suitability of the prior distributions. Fitting the models to MEG data yielded reliable posterior distributions of the parameters of each of the BNMs. Finally, model comparison revealed the BNM with “distance-dependent delays”, as the most probable to describe the generation of alpha-band networks of phase synchronization seen in MEG. These findings suggest that distance-dependent delays contribute significantly to the neocortical architecture of human alpha-band networks of phase synchronization. Hence, our study illuminates the role of inter-regional delays in generating the large-scale networks of phase synchronization that might subserve the communication between regions vital to cognition.

**Highlights:**

- Compared methods to specify delays in Biophysical Network Models (BNMs)
- BNM with “distance-dependent” conduction delays more probable than alternatives
- BNMs with biologically informed prior distributions generate dynamics seen in MEG
- Fitting BNMs yields reliable posterior distributions informed by MEG data (*N* = 75)

## 1. Introduction

Communication between brain regions is fundamental to all sensorimotor and cognitive functions (Fries (2015), Buszáki (2006), Varela et al. (2001)). Phase synchronization between neuronal oscillations from different brain regions is considered to subserve inter-regional communication by regulating the relation of spike arrival times to windows of excitability in the receiving brain region (Fries (2015), Fries (2005), Womelsdorf et al. (2007), Salazar et al. (2012)). Distinct sets of brain regions are recruited into networks of phase synchronization in tasks involving, *e.g.*, working memory (Kitzbichler et al. (2011), Palva et al. (2010)), language (Doesburg et al. (2016)), visual attention (Lobier et al. (2018), Gross et al. (2004)), and sensorimotor processing (Hirvonen et al. (2018)). Neurophysiological studies have revealed reciprocal interactions between excitatory and inhibitory neuronal populations to underlie intra-regional phase synchronization (Buzsáki (2006), Traub (1997), Gray (1994)). However, the mapping between large-scale, inter-regional networks of phase synchronization and their structural substrates, *i.e.*, the structure-function relationship, remains poorly understood.

Biophysical Network Models (BNMs) comprise models of individual brain regions linked by biologically informed patterns of anatomical connections via finite conduction delays (Woolrich & Stephan (2005)). BNMs are a powerful tool to understand the structure-function relationship pertaining to inter-regional networks of phase synchronization (Breakspear (2017)). BNMs have been used to demonstrate the influences of oscillatory activity from neuronal populations (Forrester et al. (2020)), the pattern of inter-regional anatomical connections (Finger & Bönstrup et al. (2016)), and inhibitory synaptic plasticity (Abeysuriya et al. (2018)), in generating large-scale networks of phase synchronization observed in Electroencephalography (EEG) or Magnetoencephalography (MEG) resting-state. They have also been used to relate the heterogeneity of inter-regional conduction delays to the observed (Dotson et al. (2014)) bimodal distribution in angles of inter-regional phase synchronization (Petkoski et al. (2018), Petkoski & Jirsa (2019)). However, the influence of inter-regional delays in generating the pattern of connection strengths in large-scale networks of phase synchronization observed in EEG or MEG resting-state, has not been investigated.

BNMs typically specify inter-regional delays by dividing the Euclidean distance between regions with a biologically-informed but spatially uniform conduction velocity (Abeysuriya et al. (2018), Hadida et al. (2018), Cabral et al. (2014), Nakagawa et al. (2014), Deco et al. (2009), Ghosh et al. (2008)). BNMs with “distance-dependent delays” assuming spatially uniform conduction velocity have been used to generate alpha-band (8–12 Hz) inter-regional networks of phase synchronization corresponding to those observed in MEG (Abeysuriya et al. (2018)) and EEG resting-state (Finger & Bönstrup et al. (2016)). However, a wealth of evidence from human intra-cranial EEG recordings (Trebaul et al. (2018), Lemaréchal et al. (2022)) and animal electrophysiological studies across species (Chomiak et al. (2008), Swadlow et al. (1978), Miller (1975), Swadlow (1990), Simmons & Pearlman (1983)) report spatially varying conduction velocities. Theoretical proposals have suggested that the fine temporal co-ordination in many cognitive functions requires regulating conduction velocities, to compensate for delay heterogeneity due to varying connection lengths (Seidl (2014), Pajevic et al. (2014)). Myelination of neurons can regulate conduction velocities through the linear relationship between outer axonal diameter and conduction velocity (Rushton (1951), Waxman & Bennett (1972)). Computational models incorporating activity-dependent myelination have been demonstrated to yield inter-regional connections with highly similar conduction delays, irrespective of the length of these connections (Noori et al. (2020)). Animal electrophysiological studies (Salami et al. (2003), Carr & Konishi (1990)) have also found evidence for “isochronous delays”, *i.e.*, highly similar delays, across connections, possibly as a result of activity-dependent myelination. Alternative theoretical proposals have suggested that the need for fine temporal co-ordination might be balanced by the high metabolic costs of myelinating long-distance connections (Aboitiz et al. (2003)), resulting in a combination of “distance-dependent” and “isochronous” inter-regional conduction delays. In line with this proposal, animal electrophysiological studies have found evidence for isochronous delays in ipsilateral but not contralateral connections (Chomiak et al. (2008)). However, these alternative, biologically plausible methods to specifying inter-regional delays in BNMs, have not been compared to the standard “distance-dependent delays” method.

In this study, we compared the “distance-dependent delays” method to two alternative biologically plausible methods to specifying inter-regional delays in BNMs of alpha-band (8– 12 Hz) networks of phase synchronization. We focused on alpha-band frequencies i) because they provide a basis to compare against previously proposed BNMs of phase synchronization (Abeysuriya et al. (2018), Finger & Bönstrup et al. (2016)), which also focused on alpha-band frequencies, ii) because of the clear evidence for alpha-band oscillations manifested as a spectral peak in the 8–12 Hz range both in our own MEG dataset (see Section 2.1) and in previous human electrophysiological studies (Mahjoory et al. (2020), Donoghue et al. (2020), Wang (2010)) -oscillations are a pre-requisite for phase synchronization, and iii) because of the prominent functional role of alpha-band oscillations in cognitive functions, *e.g.*, stimulus suppression, stimulus selection and top-down modulation (Palva & Palva (2007), Foxe & Snyder (2011), Klimesch (2012)). Apart from a standard BNM with “distance-dependent delays”, we defined a BNM with “isochronous delays” which assumed highly similar inter-regional delays across connections, and a BNM with “mixed delays” which assumed inter-regional delays to be a function of both the distance between regions and an isochronous or constant delay.

We followed an Approximate Bayesian Computation (ABC) workflow to compare the three BNMs. To do this, we first specified prior distributions for parameters of each of the three BNMs based on strong neurophysiological constraints derived from the aggregated animal electrophysiology literature (Tripathy et al. (2014), Tripathy et al. (2015)). In these models, prior distributions are probability distributions reflecting our existing knowledge on the values of BNM parameters, while posterior distributions are probability distributions reflecting our updated knowledge on the values of BNM parameters after accounting for evidence from MEG data. We ran Prior Predictive Checks to verify the suitability of the chosen prior distributions, and then applied Bayesian Optimisation for Likelihood Free Inference (BOLFI) (Gutmann & Corander (2016)) to separately fit the BNMs with “distance-dependent delays”, “isochronous delays”, and “mixed delays”, to an experimental MEG resting-state dataset (*N* = 75). Finally, we used ABC model comparison (Beaumont (2019)) to compare the three fitted BNMs with an independent MEG resting-state dataset (*N* = 30). The Prior Predictive Checks revealed the range of dynamics generated by the three BNMs to encompass those reflected by the phase synchronization phenomena we observed in MEG resting-state. This suggested the suitability of the prior distributions of the parameters of all three BNMs. Fitting the three BNMs to experimental MEG data with BOLFI yielded reliable posterior distributions, representing constraints on the values of BNM parameters after accounting for evidence from the MEG data. Finally, ABC model comparison revealed the BNM with “distance-dependent delays” as more probable than the other BNMs, to describe the mechanisms generating large-scale alpha-band networks of phase synchronization observed in MEG resting-state.

## 2. Materials & Methods

We used an ABC workflow to compare the “isochronous delays”, “mixed delays”, and “distance-dependent delays” methods of specifying inter-regional delays in BNMs of alpha-band networks of phase synchronization (Figure 1). First, we employed the high-dimensional ABC inference method BOLFI (Gutmann & Corander (2016)) to fit BNMs with “isochronous delays”, “mixed delays”, and “distance-dependent delays”, to an MEG resting-state dataset (*N* = 75). Then, we used ABC model comparison (Beaumont (2019)) to choose between the three fitted BNMs on an independent MEG resting-state dataset (*N* = 30). We used the ABC workflow since it provides methods to fit and compare BNMs despite their likelihood functions being intractable or mathematically difficult to formulate (Green et al. (2015), Lintusaari et al. (2017)). Further, ABC methods perform Bayesian inference (Gelman et al. (2013), van de Schoot et al. (2021), Gelman et al. (2020)), which provides a principled framework i) to combine existing knowledge from *e.g.*, animal electrophysiology with evidence from observed MEG data to estimate values of BNM parameters, and ii) to account for uncertainty in the values of BNM parameters. We express existing knowledge of BNM parameters as prior distributions while we express updated knowledge of BNM parameters, given the observed data, as posterior distributions. Marginal distributions represent the probability distributions of individual BNM parameters irrespective of the values of other BNM parameters. Conditional distributions represent the probability distributions of individual BNM parameters given the value of another BNM parameter. Joint distributions represent the probability distribution of all BNM parameters given the values of all other BNM parameters. In this paper, we refer to marginal prior and posterior distributions of BNM parameters as simply their “prior distributions” and “posterior distributions” respectively, while we refer to conditional or joint prior and posterior distributions by their entire names, *e.g.*, “conditional prior distribution”.

**Figure 1.**
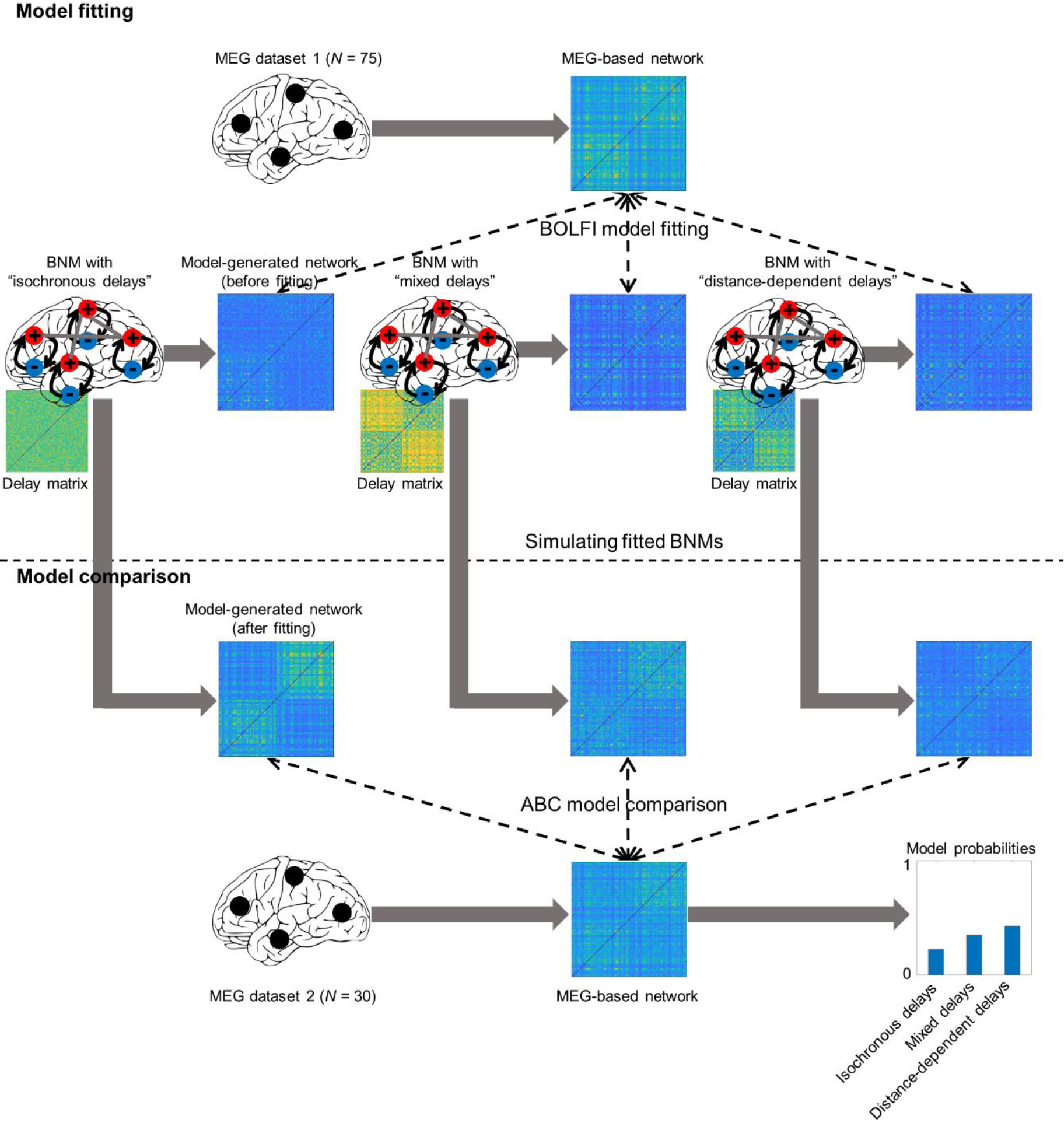
Workflow comparing strategies to specify inter-regional delays in Biophysical Network Models (BNMs) of phase synchronization. Bayesian Optimisation for Likelihood- Free Inference (BOLFI) was used to fit BNMs with “isochronous delays”, “mixed delays”, and “distance-dependent delays” to Magnetoencephalographic (MEG) resting-state data (*N* = 75), *i.e.* to determine parameter values for each of the three BNMs that would generate alpha- band inter-regional networks of phase synchronization corresponding closely to those observed in MEG resting-state. Approximate Bayesian Computation (ABC) model comparison was then used to choose between BNMs with “isochronous delays”, “mixed delays”, and “distance-dependent delays”, by comparing their alpha-band networks of phase synchronization to those observed in an independent MEG dataset (*N* = 30).

### 2.1 BNM specification

BNMs comprise models of individual brain regions linked by biologically informed patterns of anatomical connections with finite conduction delays. For BNMs implementing each of the delay specification methods, we used Wilson-Cowan (WC) oscillators to model the dynamics of individual brain regions (Wilson & Cowan (1972), Kilpatrick (2013), Cowan et al. (2016)). WC oscillators have been used to model dynamics of individual brain regions in a number of modelling studies emulating brain functional networks (Hadida et al. (2018), Hellyer et al. (2016), Heitmann et al. (2017)), including modelling studies on inter-regional networks of phase synchronization (Abeysuriya et al. (2018), Hlinka & Coombes (2012)).

The dynamics of WC oscillators arise from the interaction between excitatory and inhibitory neuronal populations, *i.e.*, the Pyramidal Inter-Neuronal Gamma (PING) model of oscillation generation (Traub et al. (1997)) and are also influenced by external inputs and the dynamics of linked oscillators. Hence, the ensemble of connected WC oscillators represented our current understanding on the generation of neuronal oscillations and inter-regional phase synchronization (Buzsáki (2006), Gray (1994)). The dynamics of oscillator *x* is given by:

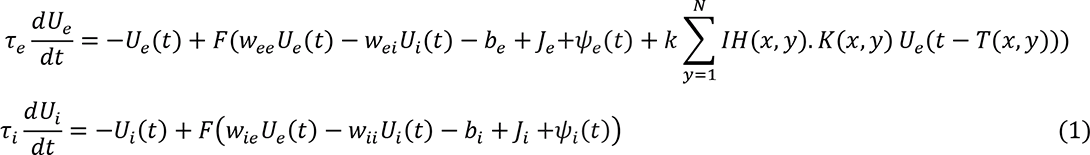

where *T* is an *N* × *N* matrix, with *T*(*x*, *y*) specifying the inter-regional conduction delay from brain region *y* to brain region *x*, in milliseconds. *N* is the number of brain regions or WC oscillators. We assumed the dynamics of each of the *N* brain regions to be governed by equation 1, in line with the assumption of identical brain regions in previous modelling studies of inter-regional phase synchronization in MEG (Abeysuriya et al. (2018), Finger & Bönstrup et al. (2016)). Further, we assumed all *N* brain regions to generate oscillatory dynamics, in agreement with the previously reported cortex-wide alpha-band spectral peaks in a large MEG resting-state dataset (*N* = 187) (Mahjoory et al. (2020)) as well as the prominent alpha-band spectral peak across regions and subjects in our own MEG dataset (*N* = 75) (Figure S1) – spectral peaks are a signature of oscillatory dynamics (Wang (2010)).

For the BNM with “distance-dependent delays”, we estimated *T*(*x*, *y*) by dividing the Euclidean distance between regions *x* and *y* (in mm) by a scalar value *v*, which was the spatially uniform conduction velocity (in metres/second) assumed by distance-dependent delays. We estimated Euclidean distance between the centroids of brain regions in MNI space. For the BNM with “isochronous delays”, we populated the upper triangular elements of *T* by sampling from a Gaussian distribution whose mean was given by a *delay* parameter (in milliseconds) and whose standard deviation was given by the product of the *delay* parameter and a *coeffvar*_*delay*_ parameter, which controlled the coefficient of variation around the mean. We constrained each of the delays to be positive using the ‘absolute’ operation and then constrained each of the delays to be integers using the ‘ceiling’ operation. Finally, we constrained the delays to be identical in both directions, *i.e.*, *T*(*x*, *y*)=*T*(*y*, *x*) for all *x* and *y* values, by copying all upper-triangular elements of *T*(*x*, *y*) to their corresponding lower-triangular elements. For the BNM with “mixed delays”, the inter-regional delays were determined both by an inter-regional distance term as well as a constant or isochronous delay term. We implemented the “mixed delays” method by first estimating the *N* × *N* velocity matrix *V*_*distance*_ implied by the distance-dependent contribution. To do this, we set all non-diagonal elements of *V*_*distance*_ to the value of the spatially uniform conduction velocity *v* assumed by distance-dependent delays. We next estimated the velocity matrix *V*_*isochronous*_ implied by the isochronous delay contribution. To do this, we divided the *N* × *N* matrix of inter-regional distances by the scalar value of *delay* parameter assumed by isochronous delays. We then combined the *V*_*distance*_ and *V*_*isochronous*_ matrices in the relative proportion specified by the *coeff*_balance_ parameter, which typically assumed values between 0 and 1. We estimated the *N* × *N* velocity matrix *V*_*mixed*_ implied by “mixed delays” by:

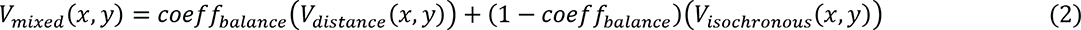

Finally, we estimated the *N* × *N* matrix *T* of “mixed delays” by an element-wise division of the *N* × *N* matrices of inter-regional distances and the *N* × *N* velocity matrix, *V*_*mixed*_. We constrained all delays to be positive using the ‘ceiling’ operation. Please refer Figure S2 for illustrations of example matrices of conduction velocities and resulting matrices of inter-regional delays for the “isochronous delays”, “mixed delays” and “distance-dependent delays” methods. 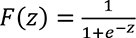 is a sigmoid function, *U*_*e*_(*t*) and *U*_*i*_(*t*) are the mean firing rates at time *t* of the excitatory and inhibitory populations respectively, *w*_*ee*_ and *w*_*ii*_ are the excitatory-excitatory and inhibitory-inhibitory connection weights respectively, *w*_*ie*_ and *w*_*ei*_ are the excitatory-inhibitory and inhibitory-excitatory connection weights, *b*_*e*_ and *b*_*i*_ correspond to the firing thresholds of excitatory and inhibitory populations, *j*_*e*_ and *j*_*i*_ are injection currents to excitatory and inhibitory populations, *ψ*_*e*_(*t*) and *ψ*_*i*_(*t*) are noise input modelled by a Gaussian process with zero mean and standard deviation given by *ψ*_*sigma*_, *τ*_*e*_ and *τ*_*i*_ are the time constants of the excitatory and inhibitory populations, and *k* is a scalar multiplier over the coupling matrix *K*, which is an *N* × *N* matrix. *K*(*x*, *y*) is the strength of the structural connection from brain region *y* to brain region *x*. *IH* is an *N* × *N* matrix that we used to selectively scale inter-hemispheric structural connections to compensate for the known under- estimation of long-distance connections by diffusion MRI-based tractography (Sotiropoulos & Zalesky (2019)). We specified the *IH* matrix by setting all elements corresponding to intra-hemispheric connections to 0, while we set all elements corresponding to inter-hemispheric connections to an identical positive value given by the *IH*_*scaling*_ parameter.

Each of the three BNMs had 11 parameters in common, *i.e.*, those parameters corresponding to the dynamics of individual brain regions and the structural connectome. In addition, the three BNMs had different sets of parameters to specify the matrix of inter-regional delays - the BNM with “distance-dependent delays” had the *v* parameter, the BNM with “isochronous delays” had the *delay* and *coeffvar*_*delay*_ parameters, while the BNM with “mixed delays” had the *v*, *delay* and *coeff*_balance_ parameters.

#### 2.1.1 Specifying strength of structural connections between WC oscillators

We specified the number and positions of brain regions as per the Destrieux brain parcellation (Destrieux et al. (2010)), whose 148 regions provided a balance between biologically detailed brain regions and computationally tractable model simulations. We specified the strengths of structural connections between WC oscillators by first estimating a 148 × 148 Destrieux atlas-based group-averaged (*N* = 57) matrix of the number of streamlines between brain regions, estimated by constrained spherical deconvolution (Smith et al. (2013)) and probabilistic tractography (Smith et al. (2012)) on pre-processed DWI images from the Human Connectome Project (van Essen et al. (2013)). The strengths of structural connections varied across seven orders of magnitude, *i.e.*, from 10^-2^ through 10^4^, and log-transformed strengths were inversely related to Euclidean distance between brain regions (Figure S3). We normalised each element in the structural connectivity matrix by its row-sum (Hlinka & Coombes (2012), Forrester (2020)). This normalisation strategy adjusts for potential tractography-induced confounds between streamline counts and sizes of brain regions. Similar strategies have been shown to improve the correspondence between diffusion MRI tractography-based structural connectivity estimates and those from retrograde tracer injections in macaque (Donahue et al. (2016)).

#### 2.1.2 Simulating the model

We simulated all three BNMs with the DDE23a integrator (Shampine & Thompson (2001)), through the Brain Dynamics Toolbox (BDT) (Heitmann et al. (2018)). We ran the model simulations for 65 seconds with a 250 Hz sampling frequency, and set Absolute Tolerance to 1 × 10^-6^ and Relative Tolerance to 1 × 10^-3^, to limit local discretisation error. We discarded data from the first 5 seconds to minimise the effect of transient dynamics. We used dynamics of only the excitatory neuronal populations for further processing since the pyramidal neurons in excitatory populations are the dominant contributors to the measured MEG signals (Lopes da Silva (2013)). The dynamics of the excitatory neuronal populations represented the mean firing rate of pyramidal neurons in these populations.

### 2.2 Prior specification

We specified prior distributions of BNM parameters as Gaussian distributions whose mean and standard deviation we set based on i) biological constraints, including values reported in the aggregated animal electrophysiology literature and human intra-cranial EEG recordings, ii) values found to be optimal in the MEG and functional Magnetic Resonance Imaging (fMRI) modelling literature on brain functional networks, and iii) ranges of values generating oscillatory dynamics - oscillations are a pre-requisite of phase synchronization.

*Prior distributions of τ*_*e*_ *and τ*_*i*_

We set the prior distribution of *τ*_*e*_, the time constant of excitatory neuronal populations, to 18.6 ± 3.6 ms (mean ± standard deviation) (Figure 2a) based on the weighted mean and pooled standard deviation of ‘layer 2/3 pyramidal neurons’ time constants in the NeuroElectro database (Tripathy et al. (2015)). We used values from ‘layer 2/3 pyramidal neurons’ since post-synaptic potentials (PSPs) from apical dendrites of supra-granular neurons are the dominant contributors to the measured MEG signal (Baillet (2017)). We set the prior distribution of *τ*_*i*_, the time constant of inhibitory neuronal populations, to 15.1 ± 4.7 ms based on the weighted mean and pooled standard deviation of time constants of different cortical inhibitory cell types in the NeuroElectro database: ‘basket cells’, ‘double bouquet cells’, ‘chandelier cells’, ‘Martinotti cells’, ‘bipolar cells’ and ‘interneurons from deep cortical layers’. We used values from diverse inhibitory cell types due to the variety of inhibitory cell types forming connections to ‘layer 2/3 pyramidal neurons’ (Markram et al. (2004)). We fixed *j*_*e*_ and *j*_*i*_, injection currents to excitatory and inhibitory populations to 0, reflecting negligible sensory and thalamic input at resting-state (Meijas et al. (2016)).

**Figure 2.**
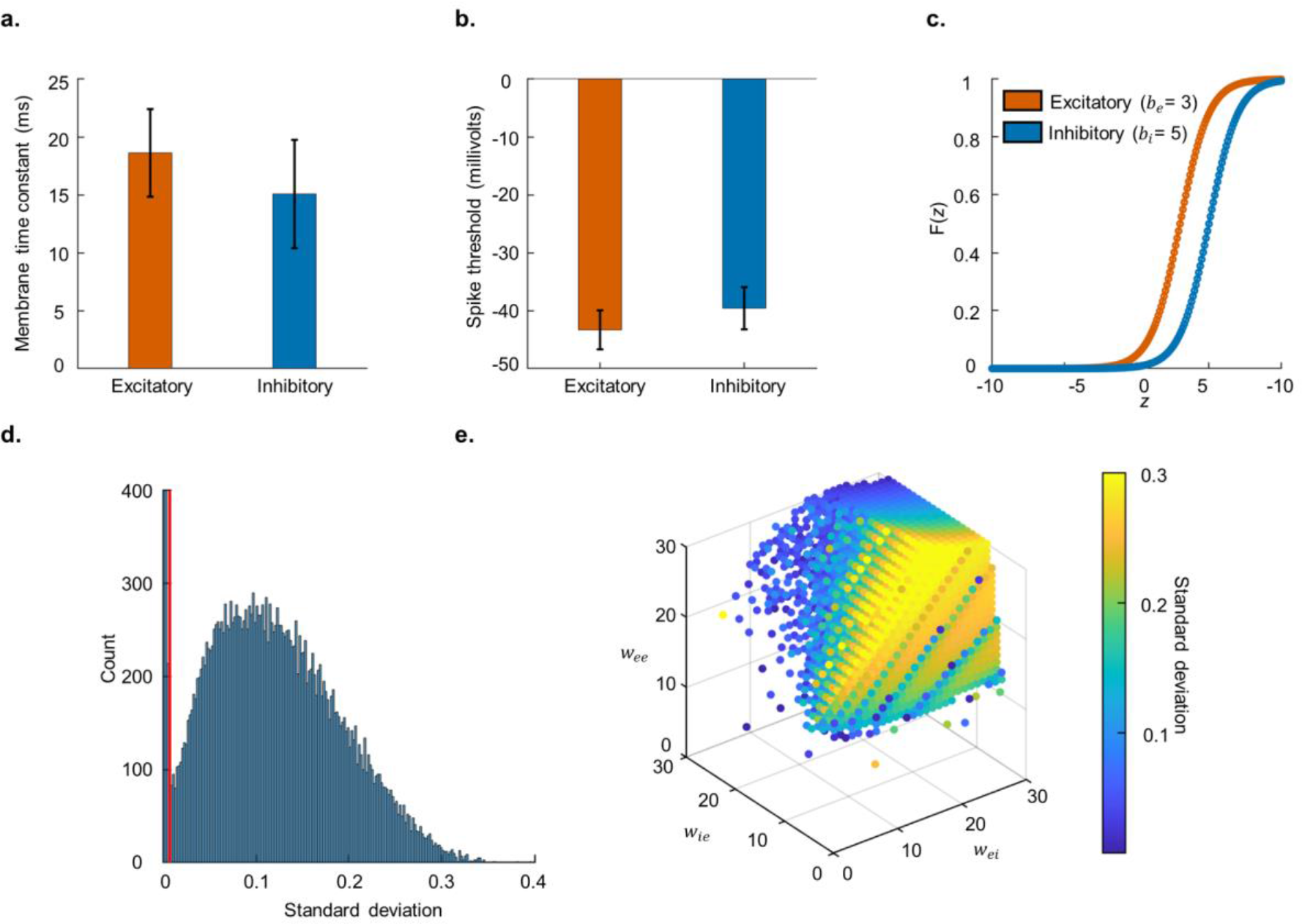
Prior specification a. Bar plot for time constants of excitatory and inhibitory neurons across multiple studies reported in NeuroElectro database. Whiskers indicate standard deviation. **b**. Bar plot for spike thresholds of excitatory and inhibitory neurons across multiple studies reported in NeuroElectro database. **c.** Logistic function curves for *b*_*e*_=3 and *b*_*i*_=5, where *b*_*e*_ and *b*_*i*_ are firing thresholds of the excitatory and inhibitory neuronal populations respectively. **d.** Histogram for standard deviation of activity from excitatory neuronal populations across multiple combinations of plausible parameter values for *w*_*ee*_, *w*_*ei*_, *w*_*ie*_ and *w*_*ii*_, which are connection strengths within and between excitatory and inhibitory neuronal populations. Vertical red line indicates standard deviation threshold of 7 × 10^-3^, to detect oscillatory dynamics. **e.** 3-D scatter plot displaying combinations of *w*_*ee*_, *w*_*ei*_ and *w*_*ie*_, generating oscillatory dynamics. Colour of dots indicates standard deviation of dynamics.

#### Prior distributions of *b*_*e*_ and *b*_*i*_

We set the prior distribution of *b*_*e*_, the firing threshold of excitatory neuronal populations, to 3 ± 1. We set the mean to a positive value since neurons fire in response to net excitation. We used a low value since the typical excitation of 20 millivolts (from -60 millivolts resting-state to -40 millivolts spike threshold) at which neurons fire, is small compared to the 100 millivolt range of the membrane potential (Kandel & Schwartz (1985)). We set the prior distribution of *b*_*i*_, the firing threshold of inhibitory neuronal populations, to 5 ± 1. We set the mean to 5 due to the higher spike thresholds of inhibitory neurons (-39.6 millivolts) compared to excitatory neurons (-43.2 millivolts) (Figure 2b), as per values in the NeuroElectro database (Tripathy et al. (2015)). Higher spike thresholds of inhibitory neurons is also in agreement with the high spike thresholds of nest basket cells, which make up a high proportion of inhibitory neurons (Wang et al. (2002)). We set the standard deviation to 1, to reflect the partial overlap in the spike thresholds of excitatory and inhibitory neurons (Figure 2b). Prior means of *b*_*e*_ and *b*_*i*_ set firing thresholds of excitatory and inhibitory populations to 3 and 5 respectively (Figure 2c).

#### Prior distributions of *w*_*ee*_, *w*_*ei*_, *w*_*ie*_ and *w*_*ii*_

We set the prior distribution of *w*_*ii*_, strength of connections within inhibitory neuronal populations, to 1 ± 0.2. These values reflected the strict biological constraint of sparse recurrent structural connectivity between inhibitory interneurons (Markram et al. (2004), Binzegger et al. (2004)). We set the prior distributions of *w*_*ee*_, *w*_*ei*_ and *w*_*ie*_, connection strengths within excitatory neuronal populations, from inhibitory to excitatory, and excitatory to inhibitory, to 20 ± 5, 18 ± 6 and 18 ± 6 respectively. These reflected ranges of parameters values generating oscillatory dynamics, as defined by a standard deviation threshold (Figure 2d–e). The higher value of the prior mean for *w*_*ee*_ compared to those of *w*_*ei*_ and *w*_*ie*_ reflected the biological constraint of dense structural connections between ‘layer 2/3 pyramidal neurons’ (Binzegger et al. (2004), Douglas et al. (1989), Douglas & Martin (2007), Jansen & Rit (1995)). The wide standard deviations for *w*_*ee*_, *w*_*ei*_ and *w*_*ie*_ reflected the uncertainty in their values due to differing reports on their relative magnitudes - anatomical studies report structural connections within excitatory populations to be much denser than those between excitatory and inhibitory populations (Binzegger et al. (2004), Douglas & Martin (2007)), while physiological studies report functional connections within excitatory populations to have similar strength to functional connections between excitatory and inhibitory populations (Seeman & Campagnola et al. (2018), Campagnola & Seeman et al. (2022)).

#### *Prior distributions of ψ*_*sigma*_, *k, and IH*_*scaling*_

We set the prior distribution of *ψ*_*sigma*_, *i.e.*, standard deviation of the noise input to excitatory and inhibitory populations, to 0.15 ± 0.05. The very low values assumed by *ψ*_*sigma*_ respected the biological constraint of negligibly small probability that a neuronal population fires solely due to noise input (Faisal et al. (2008)). Further, these settings allowed *ψ*_*sigma*_ to encompass values between 0.01 and 0.32 found to be optimal in previous MEG and fMRI modelling studies (Abeysuriya et al. (2018), Hellyer et al. (2016), Deco et al. (2009)). We set the prior distribution of *k*, the scalar multiplier over the structural connectome, to 1.5 ± 0.5. These values respected the biological constraint that extrinsic sources of excitation to brain regions are substantially weaker than intrinsic sources (Douglas & Martin (2007)). Further, these settings allowed *k* to encompass values between 1 and 3 found to be optimal in previous MEG and fMRI modelling studies (Hadida et al. (2018), Hellyer et al. (2016), Cabral et al. (2014), Deco & Jirsa (2012)). We set the prior distribution of *IH*_*scaling*_, the inter-hemispheric scaling factor over the structural connectome, to 2.5 ± 0.5. These values reflected the known underestimation of long distance connections by diffusion MRI (Sotiropoulos & Zalesky (2019)). Further, these settings allowed *IH*_*scaling*_ to encompass values between 1.5 and 3.5 found to be optimal in previous MEG modelling studies (Hadida et al. (2018)).

#### *Prior distributions of v*, *delay*, *coeffvar*_*delay*_ *and coeff*_balance_

Across the three methods, we estimated the matrix of inter-regional delays by element-wise division of the matrix of inter-regional distances by the matrix of conduction velocities.

However, each method had a different set of parameters to estimate their corresponding matrix of conduction velocities, in accordance with that method’s assumptions on spatial variation in conduction velocities (see Section 2.1). Hence, we set prior distributions for parameters specific to each of the three methods.

For the “distance-dependent delays” method, we had conduction velocity parameter *v*. We set the prior distribution of *v* to 8 ± 2 m/s. We set the mean as 8 m/s to fall within the values between 5–11 m/s reported to be optimal across several MEG and fMRI modelling studies (Abeysuriya et al. (2018), Nakagawa et al. (2014), Cabral et al. (2014), Hellyer et al. (2016), Hadida et al. (2018)). We set the standard deviation to 2 m/s, so that values from the prior distribution of *v* would encompass central tendency values between 1.1 m/s and 7.4 m/s reported across human electrophysiological (Trebaul et al. (2018), (Lemaréchal et al. (2022), Aboitiz et al. (1992)), macaque electrophysiological (Swadlow et al. (1978)) and macaque microscopy (Firmin et al. (2014)) studies. For the “isochronous delays” method, we had the mean delay parameter, *delay*, and a parameter controlling the coefficient of variation, *coeffvar*_*delay*_ . We set the prior distribution of *delay* to 10 ± 3 ms. We set the mean as 10 ms in line with the optimal “mean delay” across several MEG and fMRI modelling studies (Abeysuriya et al. (2018), Nakagawa et al. (2014), Cabral et al. (2014), Hellyer et al. (2016), Hadida et al. (2018)). We set the standard deviation to 3 ms, so that values from the prior distribution fell within the 1.5–24.9 ms range of inter-hemispheric delays reported across human and macaque electrophysiological studies (Aboitiz et al. (1992), Swadlow et al. (1978)). We set the prior distribution of *coeffvar*_*delay*_ to 0.2 ± 0.05 respectively. We chose this setting so that low values from this parameter’s prior distribution would generate nearly identical conduction delays across connections, while high values would generate sets of inter-regional delays whose variation was similar to sets of distance-dependent delays. For the “mixed delays” method, we had the *coeff*_balance_ parameter. Values between 0 and 1 indicated the relative proportion of isochronous delays and distance-dependent delays, 0 indicating fully isochronous delays. We set the prior distribution of *coeff*_balance_ to 0.5 ± 0.15, so that values from this parameter’s prior distribution generated sets of delays traversing the intermediate space between “distance-dependent” and “isochronous” sets of delays.

We refer the reader to our open dataset (Williams et al. (2023)) for time constants and spike thresholds of single studies, from which we estimated prior distributions of *τ*_*e*_, *τ*_*i*_, *b*_*e*_ and *b*_*i*_.

### 2.3 Prior Predictive Checks

Prior Predictive Checks are performed to assess the suitability of the prior distributions and the model, before proceeding to fit the model to observed data (Gelman et al. (2013), van de Schoot et al. (2021), Gelman et al. (2020)). In the Prior Predictive Checks, we used different test statistics to determine if the range of dynamics generated by the BNM encompassed those we observed in the MEG resting-state data. We ran 1,000 simulations of each of the three BNMs with parameter values drawn from their respective joint prior distributions.

Then, we estimated the values of four test statistics from the dynamics of each of the 1,000 simulations and compared the sample medians of these test statistics, to the values of those test statistics on MEG resting-state data. We estimated the following test statistics: i) median of alpha-band phase synchronization strengths between all pairs of 148 brain regions, to measure central tendency in the strengths of phase synchronization, ii) median absolute deviation (MAD) of alpha-band phase synchronization strengths between all pairs of 148 brain regions, to measure dispersion in the phase synchronization strengths, iii) mean of the Kuramoto order parameter (Kuramoto (1984), Breakspear et al. (2010)), to measure strength of zero-lag phase synchronization across the dataset, and iv) standard deviation of the Kuramoto order parameter, to measure variability in zero-lag phase synchronization across the dataset. Please see Section 2.3.2 for details.

#### 2.3.1 Processing experimental and simulated MEG data

We used eyes-open experimental MEG resting-state data from 75 subjects for ∼600 seconds, at a sampling frequency of 1000 Hz. Data was collected with a 306-channel MEG system (204 planar gradiometers and 102 magnetometers, Elekta-MEGIN Oy) at HUS BioMag laboratory, Helsinki. Ethics approval was obtained from the Ethics Committee of Helsinki University Central Hospital. The study was performed according to the guidelines in the Declaration of Helsinki. Written informed consent was obtained from each participant prior to the study. Please see Siebenhühner et al. (2020) for further details.

We used temporal Signal Space Separation (Taulu & Hari (2009)) implemented in MaxFilter to suppress extra-cranial noise, and Independent Component Analysis (ICA) in FieldTrip (Oostenveld et al. (2011)), to remove artefacts of ocular, cardiac, or muscular origin.

We estimated subject-specific forward and inverse operators to map between source space and MEG sensor space, based on individual T1-weighted anatomical MRI scans that we collected at a resolution of 1 × 1 × 1 mm with a 1.5T MRI scanner (Siemens, Germany). We processed these MRIs with FreeSurfer (http://surfer.nmr.mgh.harvard.edu/) and used the dynamic Statistical Parametric Mapping (dSPM) method (Dale et al. (2000)) implemented in MNE (Gramfort et al. (2014)) to estimate inverse operators based on subject-specific head conductivity models and cortically constrained source models. We applied fidelity weighting to these inverse operators to reduce the influence of MEG field spread (Korhonen et al. (2014)). We applied these subject-specific inverse operators to MEG sensor-level data, to reconstruct dynamics at up to 7,500 sources per hemisphere for each subject. Next, we averaged the reconstructed dynamics within each brain region in the Destrieux atlas, to obtain the representative dynamics for each of the 148 regions. We then downsampled these source collapsed datasets of each subject to 250 Hz, before bandpass filtering in the alpha frequency band (8–12 Hz) with Morlet wavelets of peak frequency = 9.83 Hz and width parameter = 5. We chose a high value for the Morlet width parameter to account for subject-wise variability in the limits of the alpha frequency band (Haegens et al. (2014)). These operations yielded 75 subject-specific alpha-band experimental MEG datasets, at the level of brain regions. From 30 of these subjects, we recorded another set of resting-state data. We used these 30 additional MEG datasets to choose between the three BNMs with ABC model comparison. Further, we recorded eyes-closed MEG resting-state data from 28 of the original cohort of 75 subjects. We used these 28 additional MEG datasets to choose between the three BNMs in eyes-closed MEG resting-state, where the compared BNMs had been fit to the original dataset of eyes-open MEG resting-state data from 75 subjects.

We generated simulated MEG data by first simulating the BNMs for 65 seconds at a sampling frequency of 250 Hz, before removing data from the first 5 seconds to remove the effect of transient dynamics. Then, we successively projected the simulated data to sensor-level with the same 75 subject-specific forward operators whose MEG data we recorded, and applied the 75 subject-specific inverse operators to the simulated sensor-level MEG data, resulting in 75 simulated source-space MEG datasets. Next, we performed the source collapsing and bandpass filtering of the simulated source-space MEG data identically as to the experimental MEG resting-state data, yielding 75 subject-specific alpha-band datasets of simulated MEG, across 148 brain regions of the Destrieux brain atlas.

#### 2.3.2 Estimating test statistics for Prior Predictive Checks

For both simulated and experimental MEG datasets, we estimated the median and median absolute deviation (MAD) of phase synchronization strengths. To do this, we first estimated subject-specific matrices of phase synchronization between all pairs of 148 brain regions from the alpha-band source-space MEG datasets of each subject. We measured phase synchronization using weighted Phase Lag Index (wPLI), which is insensitive to the confounding influence of MEG field spread on estimates of phase synchronization (Vinck et al. (2011), Siebenhühner et al. (2016), Palva et al. (2018)). We estimated wPLI as:

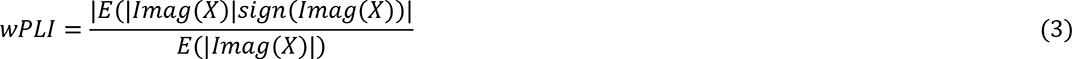

where *X* is the cross-spectrum between a pair of signals and *Imag*(*X*) is its imaginary component. We then averaged these subject-specific matrices along the subject dimension to obtain group-level matrices of phase synchronization. We estimated the median of phase synchronization strengths from the upper triangular elements of the group-level matrix of phase synchronization. We estimated the median absolute deviation (MAD) of phase synchronization strengths as the median of absolute differences between each phase synchronization strength and the median phase synchronization.

For both simulated and experimental source-space MEG datasets, we estimated the mean and standard deviation of the Kuramoto order parameter *R*, by first estimating *R* at each time *t*:

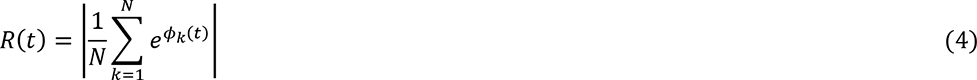

where *ϕ*_*k*_(*t*) is the instantaneous phase of the oscillator with index *k*, and *N* is the total number of oscillators. We estimated the mean and standard deviation of *R*(*t*) for the alpha-band MEG dataset of each subject and then averaged these estimates across subjects, to obtain group-level estimates of the strength and variability of zero-lag phase synchronization.

Please refer Table 1 for an overview of the test statistics we used, how we estimated them and our purpose in using them.

**Table 1.** Descriptions of each test statistic, their estimation and purpose.

| Test statistic | Estimation | Purpose |
| --- | --- | --- |
| Central tendency in strengths of inter-regional phase synchronization | Median of upper-triangular elements of group-level matrix of phase synchronization | To measure overall strength of inter-regional phase synchronization |
| Dispersion in strengths of inter-regional phase synchronization | Median Absolute Deviation (MAD) of upper-triangular elements of group-level matrix of phase synchronization | To measure overall variability of inter-regional phase synchronization |
| Strength of aggregate phase synchronization | Mean of subject-level means of Kuramoto order parameter | To measure strength of simultaneous, zero-lag phase synchronization across dataset |
| Temporal variability of aggregate phase synchronization | Mean of subject-level standard deviations of Kuramoto order parameter | To measure temporal variability of simultaneous, zero-lag phase synchronization across dataset |

### 2.4 BNM fitting

We used an ABC method, BOLFI (Bayesian Optimisation for Likelihood-Free Inference) (Gutmann & Corander (2016)) to fit each of the BNMs to experimental MEG data. We used the BOLFI implementation in the Python package, Engine for Likelihood Free Inference (ELFI) (Lintusaari et al. (2018)). We chose BOLFI to estimate BNM parameters since it is suitable for i) likelihood-free inference (LFI) settings where a model’s intractable likelihood function renders standard likelihood-based methods inapplicable (Lintusaari et al. (2017)), and ii) high-dimensional inference, *i.e.* estimating more than ∼10 model parameters - standard LFI methods such as ABC-Sequential Monte Carlo (SMC) (Sisson et al. (2007), West et al. (2021)) are only suitable to estimate a few model parameters and do not scale well to high-dimensional settings (Gutmann & Corander (2016)). BOLFI has been used to infer parameters of models in diverse fields, including genetics (Corander et al. (2017), McNally & Kallonen et al. (2019), Arnold et al. (2018)), cosmology (Leclercq (2018)), computational social science (Asikainen et al. (2020)) and cognitive science (Kangasrääsiö et al. (2019)).

While a method similar to BOLFI has been used to estimate parameters of BNMs in Systems Neuroscience (Hadida et al. (2018)), it does not perform Bayesian inference - limiting its ability to include existing *e.g.*, neurophysiological constraints on values of BNM parameters, and to account for uncertainty in the values of BNM parameters when comparing BNMs.

BOLFI estimates posterior distributions of BNM parameters using Bayes’ rule (Gelman et al. (2013)) to combine prior distributions of BNM parameters with an approximation of the BNM’s likelihood function. We employed a Gaussian Process (GP)-based surrogate model to approximate the BNM’s likelihood function. We trained the GP model with the results of multiple BNM simulations, to learn the mapping between combinations of parameter values and the corresponding discrepancies between BNM dynamics and MEG data. We used summary statistics to describe the BNM dynamics and MEG data. We used GPs due to their suitability in modelling smooth input-output relationships (Rasmussen & Williams (2006)) - we expected similar combinations of parameter values to generate similar BNM dynamics.

Previously studied BNMs have demonstrated smooth input-output relationships (Hadida et al. (2018), Perl et al. (2020)). GPs acquire their smoothness constraint from their covariance matrix. We specify the functional form of the covariance matrix with a kernel, and we use a kernel lengthscale parameter to quantify the rate of decrease in covariance with increases in values of BNM parameters. When used with BOLFI, GP surrogate models have drastically reduced the number of model simulations required to accurately estimate values of model parameters (Gutmann & Corander (2016)). Hence, we used BOLFI with GP surrogate models to fit high-dimensional BNMs of between 12 to 14 parameters in our study, to MEG data.

#### 2.4.1 BOLFI settings

We employed the following procedure and settings to apply BOLFI to estimate joint posterior distributions of each of the three BNMs. We set the prior distributions of parameters for each BNM as per the values we had specified (Section 2.2). We used the 148 × 148 group-level matrix of static phase synchronization estimated from MEG resting-state (Section 2.3.2) to represent experimentally observed dynamics, against which we compared BNM dynamics.

We chose to compare the group-level matrices of static phase synchronization estimated from the MEG data and BNM dynamics rather than corresponding descriptions of time-varying phase synchronization, due to i) the stable inter-regional patterns of phase synchronization across time reported in recent human electrophysiological studies (Nentwich et al. (2020), Mostame & Sadaghiani et al. (2021), Sadaghiani et al. (2022)), and ii) since comparing descriptions of time-varying phase synchronization returned by, *e.g.*, a HMM (Hidden Markov Model)-based method (Vidaurre et al. (2018)) would add a layer of complexity to the BNM fitting by increasing the dimensionality of the summary statistics (Lintusaari et al. (2017)) by a multiplicative factor equal to the number of hidden states and introducing problems of “state matching” between hidden states estimated from the MEG data and BNM dynamics. We simulated the BNM at 10,000 combinations of parameter values drawn from the BNM’s joint prior distribution. From the dynamics of each BNM simulation, we estimated 148 × 148 group-level matrices of phase synchronization. We chose the summary statistics to be the vector of upper-triangular elements of the 148 × 148 group-level matrices and used the Structural Similarity Index (SSI) (Wang et al. (2004)) to measure the similarity between summary statistics of the BNM dynamics and those from MEG data. We used SSI to measure similarity due to i) it simultaneously comparing mean, standard deviation and pattern of values in two input vectors in contrast to alternative measures such as, *e.g.*, Pearson Correlation which only compares the pattern of values in two input vectors, ii) its demonstrated effectiveness in comparing empirical brain functional networks to those generated by BNMs (Piccinini et al. (2021)) and generative models (Perl et al. (2020)), and ii) its reported good performance in comparing high-dimensional images in image processing applications (Ledig et al. (2017), Dong et al. (2015), Wang et al. (2004a)), which is analogous to our comparing high-dimensional vectors of phase synchronization strengths. We estimated SSI as:

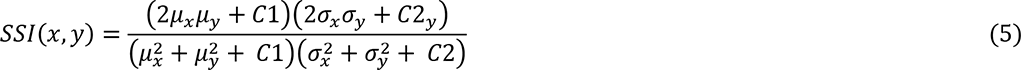

where *μ*_*x*_, *μ*_*y*_, *σ*_*x*_, *σ*_*y*_ and *σ*_*x*_*σ*_*y*_ are local means, local standard deviations and cross-covariances of the vectors *x* and *y* respectively, and *C*1 = 0.01^2^ and *C*2 = 0.03^2^. *x* and *y* were respectively the vectors of phase synchronization strengths estimated from BNM dynamics and MEG data. SSI values typically fall between 0 and 1, values close to 1 indicating highly similar vectors (Wang et al. (2004)). We expressed the discrepancy between summary statistics from MEG data and BNM dynamics as *ln*(1 − *SSI*). Hence, the discrepancy value for identical vectors would be -∞. We applied the natural logarithm to provide finer resolution at low discrepancy values (Gutmann & Corander (2016)). A single BNM simulation can exceed 24 hours, hence simulating BNMs at 10,000 combinations of parameter values in a serial manner would have prohibitively long run-time. We reduced computational run-time by exploiting the independence of BNM simulations, using an “embarrassingly parallel” paradigm on a HPC cluster to simulate BNMs at each of the 10,000 samples. We used “array jobs” to run the 10,000 simulations in 2,000 sets of 5 simulations, wherein we set the time limit for each set to 120 hours and the RAM memory limit to 30 GB. However, note that running BNM simulations in this manner only permitted training the GP model with combinations of parameter values drawn from their joint prior distributions. We did not run BNM simulations at points suggested by a Bayesian Optimisation (BO) acquisition function, *i.e*., we did not have an active learning stage in the GP training. Since BNM simulations are not independent of each other during active learning, including an active learning stage would make computational run-times prohibitively long. We used an ARD (Automatic Relevance Determination) squared exponential kernel with a constant basis function to specify the functional form for the covariance matrix of the GP surrogate model. Algorithmic complexity of fitting the GP model scales as a cube of the number of simulations (Gutmann & Corander (2016)), hence fitting the GP model to ∼10,000 points can be computationally expensive. To aid convergence, we first used the subset of data method (2,000 points) to fit the GP model and used the residual noise variance estimated from this fit as a fixed parameter when fitting the GP model to ∼10,000 points. Once the GP fitting was complete, we assessed the quality of the fit by estimating the Pearson Correlation between actual discrepancies and GP-predicted discrepancies. We also determined the relative importance of each BNM parameter in explaining the actual discrepancies, by computing *exp*^−*lengthscales*^ of the estimated ARD kernel lengthscales. Next, we estimated the posterior distributions of BNM parameters by combining the GP-based likelihood function with prior distributions of the BNM parameters. To estimate posterior distributions, we used the NUTS method (Hoffman & Gelman (2014)) to sample 1,000 points each, from 4 chains, with half these points being used for warm-up.

We set the posterior defining threshold as the minimum of the GP-based mean discrepancy function (Gutmann & Corander (2016)), and set 0.8 as the target probability, which is within the recommended range for this value (Betancourt et al. (2014)). Finally, we assessed convergence of the posterior sampling stage by checking if the effective number of samples was <100 and *R̂* <1.05, for each of the BNM parameters (Vehtari et al. (2021)). Effective number of samples indicates the number of samples from the posterior after accounting for autocorrelation between samples (Geyer et al. (2011)), while the *R̂* diagnoses “chain mixing” by comparing between-chain and within-chain estimates of model parameters - values close to 1 suggest the absence of “chain mixing”.

#### 2.4.2 Assessing sensitivity of discrepancies to values of BNM parameters

The accuracy of posterior distributions estimated by ABC methods are highly dependent on the sensitivity of the discrepancies between ‘simulated’ and ‘observed’ dynamics, to the values of the BNM parameters (Lintusaari et al. (2017), Sunnåker et al. (2013)). For BOLFI, the accuracy of the posterior distributions are also dependent on the sensitivity of the GP-predicted discrepancies to the values of BNM parameters. We used fake-data simulations to assess the sensitivity of the actual and GP-predicted discrepancies, to values of two BNM parameters *w*_*ee*_ and *w*_*ei*_ . For these fake-data simulations, we used the same BNM as specified in Section 2.1, but with “instantaneous delays” or “zero delays” - using instantaneous delays allowed us to run the BNM simulations several orders of magnitude faster since we were solving ordinary differential equations rather than delay differential equations. We first generated a reference dataset of ‘observed’ dynamics by selecting a combination of parameter values producing oscillatory dynamics. We used the following values: *w*_*ee*_ = 12.9, *w*_*ei*_ = 13.4, *w*_*ie*_ = 12.4, *w*_*ii*_ = 0.85, *b*_*e*_ = 2.85, *b*_*i*_ = 4.7, *τ*_*e*_ = 15.9, *τ*_*i*_ = 18.1, *k* = 1.6, *IH*_*scaling*_ = 2.83 and *ψ*_*sigma*_= 0.13. We simulated the BNM with these parameter values 1,000 times with ODE45 (Bogacki & Shampine (1996)), other settings being identical to that specified in Section 2.1.2. For each of the 1,000 simulations, we generated group-level matrices of phase synchronization, then averaged across these 1,000 group-level matrices to generate the reference group-level matrix of phase synchronization. Next, we generated datasets of ‘simulated’ dynamics by running 20 BNM simulations at every point in the 100 × 100 grid defined by every pairwise combination of *w*_*ee*_ and *w*_*ei*_ values. We varied *w*_*ee*_ and *w*_*ei*_ across 100 equally spaced points from 10 to 30 and from 6 to 30 respectively. We fixed values of all other BNM parameters to the same value as for the reference dataset. From the datasets of ‘simulated’ dynamics, we generated 20 group-level matrices of phase synchronization for every point in the 100 × 100 grid, and averaged across these 20 repetitions to obtain a single group-level matrix at each point in the 100 × 100 grid. Then, we estimated discrepancies between the reference ‘observed’ summary statistics and ‘simulated’ summary statistics at every point on the 100 × 100 grid. We then determined if the discrepancy surface reached a global minimum at the point on the grid representing the combination of true values of *w*_*ee*_ and *w*_*ei*_ . Further, we estimated a GP surrogate model relating the BNM parameter values to the corresponding discrepancies. We determined if the surface of GP-predicted discrepancies reached a global minimum at the point on the grid representing the combination of true values of *w*_*ee*_ and *w*_*ei*_. These investigations revealed if the actual and GP-predicted discrepancies were sensitive to the values of two BNM parameters, *w*_*ee*_ and *w*_*ei*_.

### 2.5 BNM evaluation

We evaluated the three fitted BNMs by comparing the posterior distributions of each of the BNM parameters to their respective prior distributions. Comparing the posterior distributions of BNM parameters to their prior distributions revealed additional constraints on the values of these parameters learnt from the MEG data, through BOLFI model fitting. Further, we ran Posterior Predictive Checks to assess the similarity between dynamics from the fitted BNMs and those reflected by the phase synchronization phenomena in the observed MEG data.

#### 2.5.1 Posterior Predictive Checks

We used Posterior Predictive Checks (Gelman et al. (2013), Gelman et al. (2020), van de Schoot et al. (2021)) to determine if the dynamics generated by the three fitted BNMs correspond to those reflected by the phase synchronization phenomena in the MEG data. We ran 1,000 simulations of each of the three BNMs with parameter values drawn from their respective joint posterior distributions. Just as for the Prior Predictive Checks (Section 2.3), we then estimated the values of four test statistics from the dynamics of each of the 1,000 BNM simulations and compared the sample medians of these test statistics to the values of those test statistics on experimental MEG resting-state data. We used the same set of test statistics as for the Prior Predictive Checks: i) median of alpha-band phase synchronization strengths between all pairs of 148 brain regions, ii) median absolute deviation (MAD) of alpha-band phase synchronization strengths between all pairs of 148 brain regions, iii) mean of Kuramoto order parameter, and iv) standard deviation of Kuramoto order parameter (see Table 1 for details).

### 2.6 BNM comparison

We used standard ABC model comparison to compare the fitted BNMs with “isochronous delays”, “mixed delays”, and “distance-dependent delays”. We simulated the three BNMs, each with 1,000 sets of parameter values drawn from their respective joint posterior distributions. We simulated the BNMs at samples from their joint posterior distributions rather than their joint prior distributions since the posteriors represent probable values of BNM parameters after combining information from both previous neurophysiology experiments and our own MEG data. In contrast, the priors represent probable values of BNM parameters based only on information from previous neurophysiological experiments. Hence, the posteriors are more likely than the priors to reflect the ground-truth values of the BNM parameters. It follows from this that comparing the BNMs with samples from their respective posterior distributions enables isolating the influence of delays-related BNM parameters by reducing the potentially confounding effect of inaccurate estimates of other BNM parameters on the model comparison. For each of the three BNMs, we estimated discrepancies between dynamics from each of the 1,000 simulations to dynamics from an independent dataset of MEG resting-state data (*N* = 30). We estimated discrepancy as *ln*(1 − *SSI*), identical to the original BNM fitting (Section 2.4.1). SSI is the Structural Similarity Index between the vectors of inter-regional phase synchronization strengths from BNM dynamics and MEG data. We estimated probability of each BNM by the relative acceptance rate of discrepancies associated with that BNM, with respect to a specified minimum discrepancy (Beaumont (2019)). We estimated model probabilities for a range of minimum discrepancies between -1 and 0, where -1 corresponded to a conservative threshold accepting very few discrepancy values across BNMs while 0 corresponded to a liberal threshold. We then chose between the three BNMs based on the model probabilities across a range of discrepancy thresholds.

We refer the reader to our GitHub repository for the Python and MATLAB code, and SLURM scripts (https://github.com/nitinwilliams/eeg_meg_analysis/tree/master/MEGMOD), that we used to simulate, fit and compare the BNMs. Within the GitHub repository, please check file_descriptions.txt for names of files implementing 1.) MATLAB functions to simulate each of the three BNMs – we called each of these functions via “array jobs” implemented in SLURM scripts (to be run on HPC resources), which we also make available, 2.) MATLAB code to estimate the input set of parameter values and output set of discrepancies for BOLFI model fitting, for each BNM, 3.) Python code to use the ELFI toolkit to fit each of the BNMs to MEG resting-state data with BOLFI, 4.) MATLAB code to generate the set of posterior distributions returned by BOLFI in the correct order and scale, 5.) MATLAB functions to simulate each of the three BNMs with samples from their posterior distributions – we called each of these functions via “array jobs” implemented in SLURM scripts (to be run on HPC resources), which we also make available, and 6.) MATLAB code implementing ABC model comparison to compare the three fitted BNMs.

## 3. Results

We compared the “isochronous delays”, “mixed delays”, and “distance-dependent delays” methods of specifying inter-regional delays in BNMs of alpha-band networks of phase synchronization. We specified BNMs implementing each of the three methods and then used an ABC workflow to adjudicate between them. The steps we followed were: i) we employed constraints from previous human and animal electrophysiological studies as well as the MEG and fMRI modelling literature, to specify prior distributions for parameters of each BNM, ii) we used Prior Predictive Checks to determine whether each of the BNMs, constrained by their prior distributions, generated dynamics encompassing those reflected by the phase synchronization phenomena in the MEG data, iii) we used fake-data simulations to verify that the estimated discrepancies between BNM dynamics and MEG data, were sensitive to the values of two BNM parameters, iv) we applied BOLFI to fit each of three BNMs to MEG resting-state data (*N* = 75), yielding posterior distributions of their parameters, v) we employed Posterior Predictive Checks to verify that the fitted BNMs generated dynamics corresponding closely to those observed in the MEG dataset they were trained on, and vi) we applied ABC model comparison to determine which of the three fitted BNMs generated alpha-band networks of phase synchronization most similar to those observed in an independent MEG resting-state dataset (*N* = 30).

### 3.1 Prior specification

We combined the prior distribution of BNM parameters with an approximation of the BNM likelihood function to estimate the posterior distributions of BNM parameters. Hence, using biologically plausible, well-motivated prior distributions was important to accurately estimating the posterior distributions of BNM parameters. We set prior distributions of BNM parameters based on biological constraints, parameter values found to be optimal in the MEG and fMRI modelling literature, and ranges of values generating oscillatory dynamics. We set the priors to be Gaussian distributed and list their means and standard deviations below, along with brief rationales for choosing these values (Table 2). We refer the reader to Materials & Methods, Section 2.2 (see Figure 2) for a detailed description of the prior specification.

**Table 2.** Means and standard deviations of prior distributions for each of the BNM parameters, along with brief rationales for choosing the specified values.

| Parameter<br>(description) | Mean ± SD<br>(units) | Rationale |
| --- | --- | --- |
| $w_{ee}$<br>(Connection strength<br>within excitatory<br>neuronal<br>populations) | $20 \pm 5$<br>(a.u.) | i) Dense recurrent structural connectivity between ‘layer 2/3 pyramidal neurons’ (Binzegger et al. (2004), Douglas et al. (1989), Douglas & Martin (2007), Jansen & Rit (1995))<br>ii) Similar strength of functional connections to those between excitatory and inhibitory neuronal populations *i.e.*, $w_{ei}$ and $w_{ie}$ (Seeman & Campagnola et al. (2018), Campagnola & Seeman et al. (2022))<br>iii) Encompasses range of values generating oscillatory dynamics |
| $w_{ei}$<br>(Connection strength<br>from inhibitory to<br>excitatory neuronal<br>populations) | $18 \pm 6$<br>(a.u.) | i) Weaker strength of structural connections compared to dense connectivity within excitatory neurons (Binzegger et al. (2004), Douglas & Martin (2007))<br>ii) Similar strength of functional connections to those within excitatory neurons and from excitatory to inhibitory neurons *i.e.*, $w_{ee}$ and $w_{ie}$ (Seeman & Campagnola et al. (2018), Campagnola & Seeman et al. (2022))<br>iii) Encompasses range of values generating oscillatory dynamics |
| $w_{ie}$<br>(Connection strength from excitatory to inhibitory neuronal populations) | $18 \pm 6$<br>(a.u.) | i) Weaker strength of structural connections compared to dense connectivity within excitatory neurons (Binzegger et al. (2004), Douglas & Martin (2007))<br>ii) Similar strength of functional connections to those within excitatory neurons and from excitatory to inhibitory neurons *i.e.*, $w_{ee}$ and $w_{ie}$ (Seeman & Campagnola et al. (2018), Campagnola & Seeman et al. (2022))<br>iii) Encompasses range of values generating oscillatory dynamics |
| $w_{ii}$<br>(Connection strength within inhibitory neuronal populations) | $1 \pm 0.2$<br>(a.u.) | Sparse recurrent structural connectivity between inhibitory neurons (Markram et al. (2004), Binzegger et al. (2004)) |
| $b_e$<br>(Firing threshold of excitatory neuronal populations) | $3 \pm 1$ (a.u.) | Low positive value since neurons fire in response to small, net excitation |
| $b_i$<br>(Firing threshold of inhibitory neuronal populations) | $5 \pm 1$ (a.u.) | i) Low positive value since neurons fire in response to small net excitation<br>ii) Spike thresholds of inhibitory neurons are higher than spike thresholds of excitatory neurons, across all studies reported in NeuroElectro database |
| $\tau_e$<br>(Time constant of excitatory neuronal populations) | $18.6 \pm 3.6$<br>(ms) | Time constants of ‘layer 2/3 pyramidal neurons’ across studies in NeuroElectro database |
| $\tau_i$<br>(Time constant of inhibitory neuronal populations) | $15.1 \pm 4.7$<br>(ms) | Time constants of ‘basket cells’, ‘double bouquet cells’, ‘chandelier cells’, ‘Martinotti cells’, ‘bipolar cells’ and ‘interneurons from deep cortical layers’ reported across all studies in NeuroElectro database |
| $k$ | $1.5 \pm 0.5$<br>(a.u.) | i) Low positive value since extrinsic input to excitatory population much lower than intrinsic input (Douglas & Martin (2007)) |
| (Scalar multiplier over structural connectome) |  | ii) Encompassing range of values reported in MEG and fMRI modelling literature (Hadida et al. (2018), Hellyer et al. (2016), Cabral et al. (2014), Deco & Jirsa (2012)) |
| $IH_{scaling}$<br>(Inter-hemispheric scaling factor over structural connectome) | $2.5 \pm 0.5$<br>(a.u.) | i) Known under-estimation of long-distance connections by diffusion MRI (Sotiropoulos & Zalesky (2019))<br>ii) Encompassing range of values reported in MEG modelling literature (Hadida et al. (2018)) |
| $\psi_{sigma}$<br>(Standard deviation of noise to excitatory and inhibitory populations) | $0.15 \pm 0.05$<br>(a.u.) | i) Positive value, much lower than firing thresholds of excitatory ($b_e$) and inhibitory ($b_i$) populations, since negligible probability of population firing due to noise input (Faisal et al. (2008))<br>ii) Encompassing range of values reported in MEG and fMRI modelling literature (Abeyesuriya et al. (2018), Hellyer et al. (2016), Deco et al. (2009)) |
| $v$<br>(Conduction velocity) | $8 \pm 2$ (m/s) | i) Within range of conduction velocities and corresponding axonal diameters reported in human and animal neuroanatomical and neurophysiological studies (Trebaul et al. (2018), Lemaréchal et al. (2022), Swadlow et al. (1978)), Aboitiz et al. (1992), Firmin et al. (2014))<br>ii) Encompassing range of values reported in MEG and fMRI modelling literature (Abeyesuriya et al. (2018), Nakagawa et al. (2014), Cabral et al. (2014), Hellyer et al. (2016), Hadida et al. (2018)) |
| $delay$<br>(Mean conduction delay) | $10 \pm 3$ (ms) | i) Within range of inter-hemispheric delays reported in human and animal electrophysiological studies (Aboitiz et al. (1992), Swadlow et al. (1978))<br>ii) Encompassing range of values reported in MEG and fMRI modelling literature (Abeyesuriya et al. (2018), Nakagawa et al. (2014), Cabral et al. (2014), Hellyer et al. (2016), Hadida et al. (2018)) |
| $coeffvar_{delay}$<br>(Coefficient of variation in conduction delays) | $0.2 \pm 0.05$<br>(a.u.) | i) Low values of parameter would generate sets of nearly identical inter-regional delays<br>ii) High values of parameter would generate sets of inter-regional delays with similar variation to sets of “distance-dependent delays” |
| $coeff_{balance}$ | $0.5 \pm 0.15$<br>(a.u.) | Specified to enable the “mixed delays” method to traverse the intermediate space between the |
| (Coefficient of balance between “distance-dependent” and “isochronous” delays) |  | “distance-dependent delays” and “isochronous delays” methods |

### 3.2 BNMs simulated at prior means generate alpha-band dynamics

A pre-requisite for alpha-band phase synchronization is alpha-band oscillatory dynamics from individual brain regions. Hence, we investigated if the BNMs generated oscillatory dynamics at alpha-band frequencies. To do so, we ran 10 second simulations of BNMs with “isochronous delays”, “mixed delays”, and “distance-dependent delays” at their respective prior means. Then, we determined the peak frequencies of their dynamics - oscillations manifest as peaks in frequency spectra. We found that each of the three BNMs generated oscillatory dynamics (Figure 3a–c) with mean amplitude of 0.15 and mean standard deviation of 0.08 across brain regions. These oscillatory dynamics had spectral peaks in alpha-band (Figure 3d–f), with peak frequencies of 12.9 ± 0.07 Hz (mean±standard deviation), 12.9 ± 0.1 Hz and 12.8 ± 0.14 Hz for BNMs with “isochronous delays”, “mixed delays” and “distance-dependent delays” respectively, across regions (Figure 3g–i). The mean peak frequencies of all BNMs fell within the 10.3 Hz ± 2.8 Hz distribution of alpha-band peak frequencies reported in experimental MEG data (Haegens et al. (2014)). Hence, the three BNMs simulated at their respective prior means generated alpha-band oscillations, fulfilling a pre- requisite to investigate large-scale, alpha-band networks of phase synchronization.

**Figure 3.**
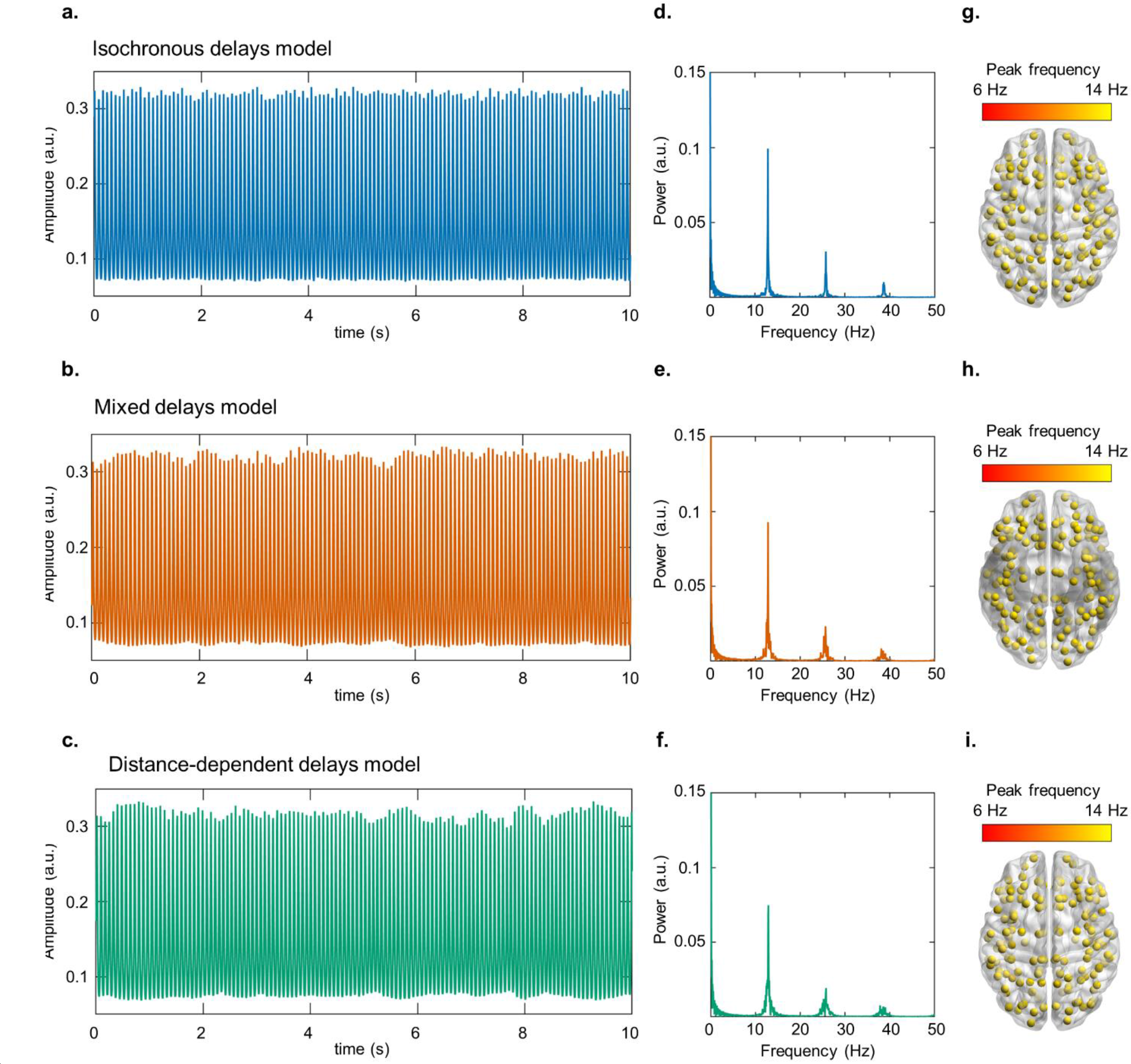
BNMs simulated at prior means generate alpha-band oscillatory dynamics. a-c. 10 s time course of dynamics from ‘left fronto-marginal gyrus and sulcus’ of BNMs with “isochronous delays”, “mixed delays”, and “distance-dependent delays” respectively. **d-f.** Frequency spectra of dynamics from ‘left fronto-marginal gyrus and sulcus’, of all three BNMs. **g-i.** Alpha-band peak frequencies of each region, of all three BNMs, in dorsal view. Plots on brain surface were visualised with BrainNet Viewer (Xia et al. (2013)).

### 3.3 BNM dynamics encompass those observed in MEG data

The BOLFI fitting method assumes the suitability of the prior distributions of the BNM parameters and that the BNMs are not mis-specified. Hence, we performed Prior Predictive Checks to assess the ability of the BNMs, constrained by their prior distributions, to generate the phase synchronization phenomena observed in MEG resting-state (Gelman et al. (2020), van de Schoot et al. (2021)). In addition, the Prior Predictive Checks allowed us to assess the similarity of the phase synchronization phenomena generated by the three BNMs, when these BNMs were constrained by their respective prior distributions. We performed the Prior Predictive Checks by comparing the sample medians of four test statistics that we estimated from 1,000 simulations of each of the BNMs, against the value of those same test statistics estimated on the experimental MEG dataset (*N* = 75). We simulated the three BNMs with parameter values drawn from their joint prior distributions. As the test statistics, we used the median and median absolute deviation (MAD) of phase synchronization strengths between all region pairs, to measure their central tendency and dispersion respectively. We also estimated the mean and standard deviation of the Kuramoto order parameter, to measure overall strength and variability of zero-lag phase synchronization respectively (see Section 2.3.2 and Table 1 for details of each test statistic). We found that the values of each of the four test statistics estimated on the MEG dataset lay within the range of values of those test statistics estimated from the dynamics of each of the three BNMs (Figure 4a–l). The dispersion in strengths of inter-regional phase synchronization estimated on the MEG dataset was 0.02, which was close to the median values of 0.03, 0.02 and 0.02 for this test statistic, for the “isochronous delays”, “mixed delays”, and “distance-dependent delays” methods, respectively (Figure 4d–f). However, the central tendency of 0.09 for the strengths of inter-regional phase synchronization estimated on the MEG dataset was distant from the median values of 0.7, 0.78 and 0.79 for this test statistic, for the three methods, respectively (Figure 4a–c). The mean and standard deviation of the Kuramoto order parameter had bimodal distributions for the sets of values estimated from dynamics of each of the three BNMs.

**Figure 4.**
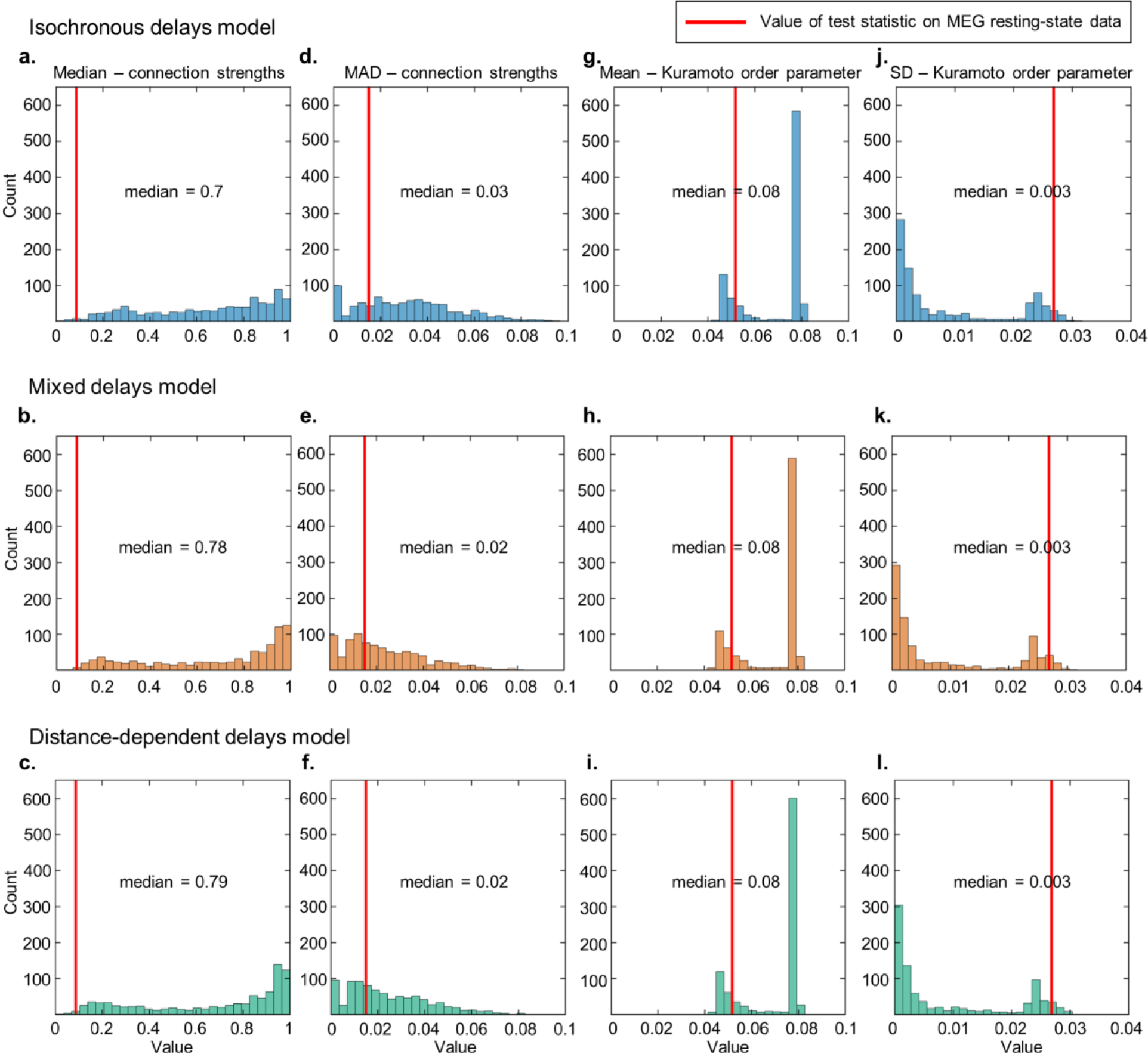
BNM dynamics encompass those observed in MEG data. a-c. Histograms of median of alpha-band phase synchronization strengths from multiple BNM simulations, where parameter values were drawn from joint prior distributions of BNMs with “isochronous delays”, “mixed delays”, and “distance-dependent delays” respectively. **d-f.** Histograms of median absolute deviation (MAD) of alpha-band phase synchronization strengths from multiple BNM simulations, of the three BNMs respectively. **g-i.** Histograms of mean of Kuramoto order parameter from multiple BNM simulations, of the three BNMs respectively. **j-l.** Histograms of standard deviation (SD) of Kuramoto order parameter from multiple BNM simulations, of the three BNMs respectively. For all panels, the red line indicates the corresponding value of that test statistic estimated from the MEG dataset.

Kuramoto mean and standard deviation close to 0.08 and 0 respectively, reflected parameter combinations for which the BNMs did not generate oscillatory dynamics while values close to 0.05 and 0.025 respectively, reflected parameter combinations for which the BNMs generated oscillatory dynamics. We found the values of 0.05 and 0.03 respectively, of these test statistics on the MEG dataset, to be close to their values for cases when the BNMs generated oscillatory dynamics (Figure 4g–l). The Prior Predictive Checks suggest that the three BNMs generate dynamics encompassing those reflected by the phase synchronization phenomena in MEG resting-state data. This suggests the suitability of the prior distributions of the BNM parameters and that the BNMs are not mis-specified, and hence can be fit to the MEG data with the BOLFI method. In addition, the correspondence between the three BNMs in the values of each of the test statistics (Figure 4a–l), suggested that each of the BNMs, constrained by their prior distributions, generate similar phase synchronization phenomena.

### 3.4 Discrepancies between BNM dynamics and MEG data are sensitive to values of BNM parameters

BOLFI returning accurate posterior distributions is highly dependent on whether the estimated discrepancies between BNM dynamics and MEG data are sensitive to values of the BNM parameters (Lintusaari et al. (2017), Sunnåker et al. (2013)). In the asymptotic case, BOLFI assumes the surface of discrepancies between summary statistics of BNM dynamics and MEG data to have a global minimum at the combination of true parameter values. We used fake-data simulations to assess this for two BNM parameters, *w*_*ee*_ and *w*_*ei*_. To do so, we first generated the ‘observed’ summary statistics as the vector of phase synchronization strengths between all region pairs, averaged across 1,000 BNM simulations. We ran the BNM simulations with a pre-chosen set of parameters values, with *w*_*ee*_ = 12.9 and *w*_*ei*_ = 13.4. Then, we generated ‘simulated’ summary statistics as the vector of phase synchronization strengths, averaged across 20 BNM simulations. We generated ‘simulated’ summary statistics at every point on a 100 ×100 grid defined by every pair of *w*_*ee*_ and *w*_*ei*_ values, where we varied *w*_*ee*_ from 10–30 and *w*_*ei*_ from 6–30. We fixed values of other BNM parameters to the same values used to generate the ‘observed’ summary statistics. Finally, we estimated the discrepancies as *ln*(1 − *SSI*) between the reference ‘observed’ summary statistics and the ‘simulated’ summary statistics at each point on the 100 ×100 grid. SSI is the Structural Similarity Index. We also estimated a set of Gaussian Process (GP)-predicted discrepancies from a GP model trained with the set of actual discrepancies and corresponding BNM parameter values. We found that the surface of actual discrepancies reached a global minimum at the combination of the true parameter values, *i.e.*, *w*_*ee*_ =12.9, *w*_*ei*_ =13.4 (Figure 5). In addition, we found low discrepancies at points on the grid corresponding to high values of *w*_*ee*_ and *w*_*ei*_, but these values were higher than the discrepancy value at the combination of true parameter values. For example, we estimated a discrepancy of -2.48 at the combination of true values (*w*_*ee*_ =12.9, *w*_*ei*_ =13.4), while we estimated a discrepancy of -2.06 at *w*_*ee*_ =27.6, *w*_*ei*_ =24.2. Notably, the surface of GP-predicted discrepancies also reached a global minimum at the combination of the true parameter values (Figure S4). These results demonstrate the sensitivity of the discrepancies to the values of *w*_*ee*_ and *w*_*ei*_, suggesting that BOLFI can return accurate posterior distributions of at least these two BNM parameters.

**Figure 5.**
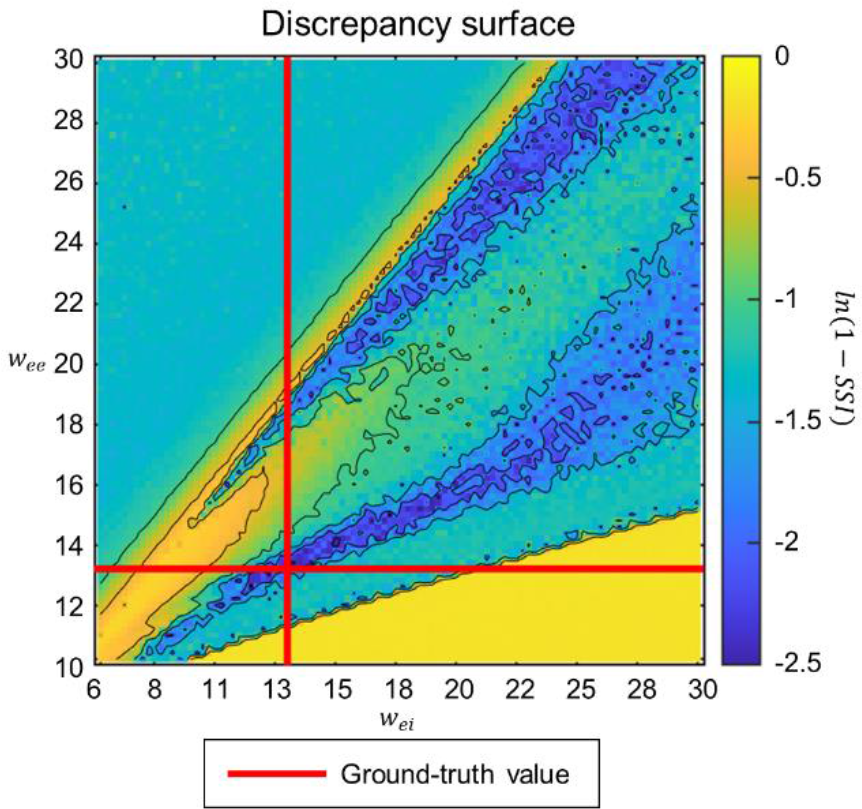
Discrepancies are sensitive to values of BNM parameters. 100 ×100 grid of discrepancies between summary statistics of BNM dynamics at every pair of *w*_*ee*_ and *w*_*ei*_ values, and BNM dynamics at *w*_*ee*_ =12.9, *w*_*ei*_=13.4. *w*_*ee*_ is the strength of connections within excitatory neuronal populations, *w*_*ei*_ is the strength of connections from inhibitory to excitatory neuronal populations. Red lines indicate ground-truth values of *w*_*ee*_ and *w*_*ei*_. Discrepancies were measured by *ln*(1 − *SSI*)). SSI is the Structural Similarity Index.

### 3.5 BOLFI yields BNM parameter estimates informed by MEG data

The behaviour of BNMs is highly dependent on the parameter values with which they are simulated. So, we first constrained values of the parameters of each of the three BNMs with MEG data, before proceeding to compare the three BNMs. To do so, we applied the high-dimensional inference method BOLFI (Gutmann & Corander (2016)), to fit each of the BNMs to MEG resting-state data (*N* = 75). BOLFI uses standard Bayesian inference to combine the prior distributions of BNM parameters with an approximation of the BNM’s likelihood function, to estimate posterior distributions of BNM parameters. BNMs typically have intractable likelihood functions, so BOLFI approximates these with Gaussian Process (GP) models trained on parameters values of multiple BNM simulations and the corresponding discrepancies between BNM dynamics and MEG data. We ran 10,000 simulations of each of the three BNMs and trained GPs parameterised with ARD squared exponential kernels, on values of the BNM parameters and the corresponding discrepancies. We estimated discrepancies as *ln*(1 − *SSI*)) between the vectors of inter-regional phase synchronization strengths estimated from BNM dynamics and MEG data. SSI is the Structural Similarity Index (Wang et al. (2004)).

The multiple BNM simulations yielded 9004, 9063, and 9093 completed simulations of BNMs with “isochronous delays”, “mixed delays”, and “distance-dependent delays” respectively, the others exceeding the time limit or crossing the memory limit. These “out of memory” errors likely reflect the excessive memory demand due to very small step sizes taken by the solver when dealing with discontinuities in the solution of the system of differential equations representing each BNM. For the completed simulations, we found Pearson Correlations between actual and GP-predicted discrepancies of 0.58, 0.67 and 0.67, for BNMs with “isochronous delays”, “mixed delays”, and “distance-dependent delays” respectively (Figure S5a–c). These close correspondences suggested the GP-based models of each BNM to suitably approximate their likelihood functions. For all three BNMs, we found that parameters governing dynamics of individual brain regions had a strong influence on predicting the discrepancies between BNM dynamics and MEG data (Figure S5d–f). In particular, the strength of connections within excitatory neuronal populations (*w*_*ee*_), between excitatory and inhibitory populations (*w*_*ei*_ and *w*_*ie*_), and the firing thresholds of excitatory (*b*_*e*_) and inhibitory populations (*b*_*i*_), had a strong influence. The influence of these parameters is consistent with neurophysiological studies on, *e.g.*, the role of reciprocal interaction between excitatory and inhibitory populations, in generating the oscillatory dynamics necessary for inter-regional phase synchronization (Buzsáki (2006), Traub (1997)). We also found that the parameter controlling the strength of inter-regional anatomical connections (*k*) had an influence on predicting the discrepancies between BNM dynamics and MEG data. This is also consistent with understanding on the role of these connections in promoting inter-regional phase synchronization (Gray (1994)).

BOLFI yielded reliable posterior distributions of parameters of all three BNMs. *R̂* values were lower than 1.05 for all parameters and effective numbers of samples exceeded 100 (Vehtari et al. (2021)) for all but one parameter, *i.e.*, *b*_*i*_ in the BNM with “isochronous delays” which had 91 effective samples. For all three BNMs, the mass of the posterior distributions of *b*_*e*_ shifted toward lower values compared to their prior distributions while the mass of the posterior distributions of *b*_*i*_ shifted toward higher values (Figure 6a–c). For the BNM with “isochronous delays” for example, prior means for *b*_*e*_ and *b*_*i*_ were 3 and 5 respectively, while their posterior means were 2.7 and 5.4 (Figure 6a). These posterior distributions of *b*_*e*_ and *b*_*i*_ across BNMs, are in agreement with neurophysiological constraints that spike thresholds of inhibitory neurons are higher than spike thresholds of excitatory neurons (see Figure 2b and Section 2.2 on Prior Specification). For all three BNMs, we also observed the mass of the posterior distributions of *τ*_*i*_, time constant of inhibitory neuronal populations, to shift toward higher values compared to their prior distributions (Figure 6a–c). For the BNM with “isochronous delays” for example, prior mean for *τ*_*i*_ was 15.1 ms while its posterior mean was 16.4 ms (Figure 6a). For BNMs with “mixed delays” and “distance-dependent delays”, mass of the posterior distributions of *w*_*ee*_ and *w*_*ie*_ shifted toward lower values (Figure 6b–c). For the BNM with the “distance-dependent delays” for example, prior means for *w*_*ee*_ and *w*_*ie*_ were 20 and 18 respectively, while their posterior means were 18.5 and 14.5 (Figure 6c). Notably, the lower values of *w*_*ee*_ were in better agreement with empirical estimates of functional connectivity within excitatory neuronal populations (Seeman et al. (2018), Campagnola et al. (2022)) than corresponding estimates of structural connectivity (Jansen & Rit (1995), Douglas & Martin (2007), Douglas et al. (1989), Binzegger et al. (2004)). We also inspected posterior distributions of BNM parameters relating to inter-regional delays. For BNMs with “isochronous delays” and “mixed delays”, the posterior means of the delay parameter *delay* were shifted to 9.5 ms and 9.6 ms respectively, from their prior means of 10 ms (Figure 6a–b). Taken together, we found that applying BOLFI yielded reliably estimated posterior distributions of parameters of the three BNMs, which were in agreement with neurophysiological results. Hence, we could use these BNMs, constrained by MEG data, to choose between the three methods to specify inter-regional delays.

**Figure 6.**
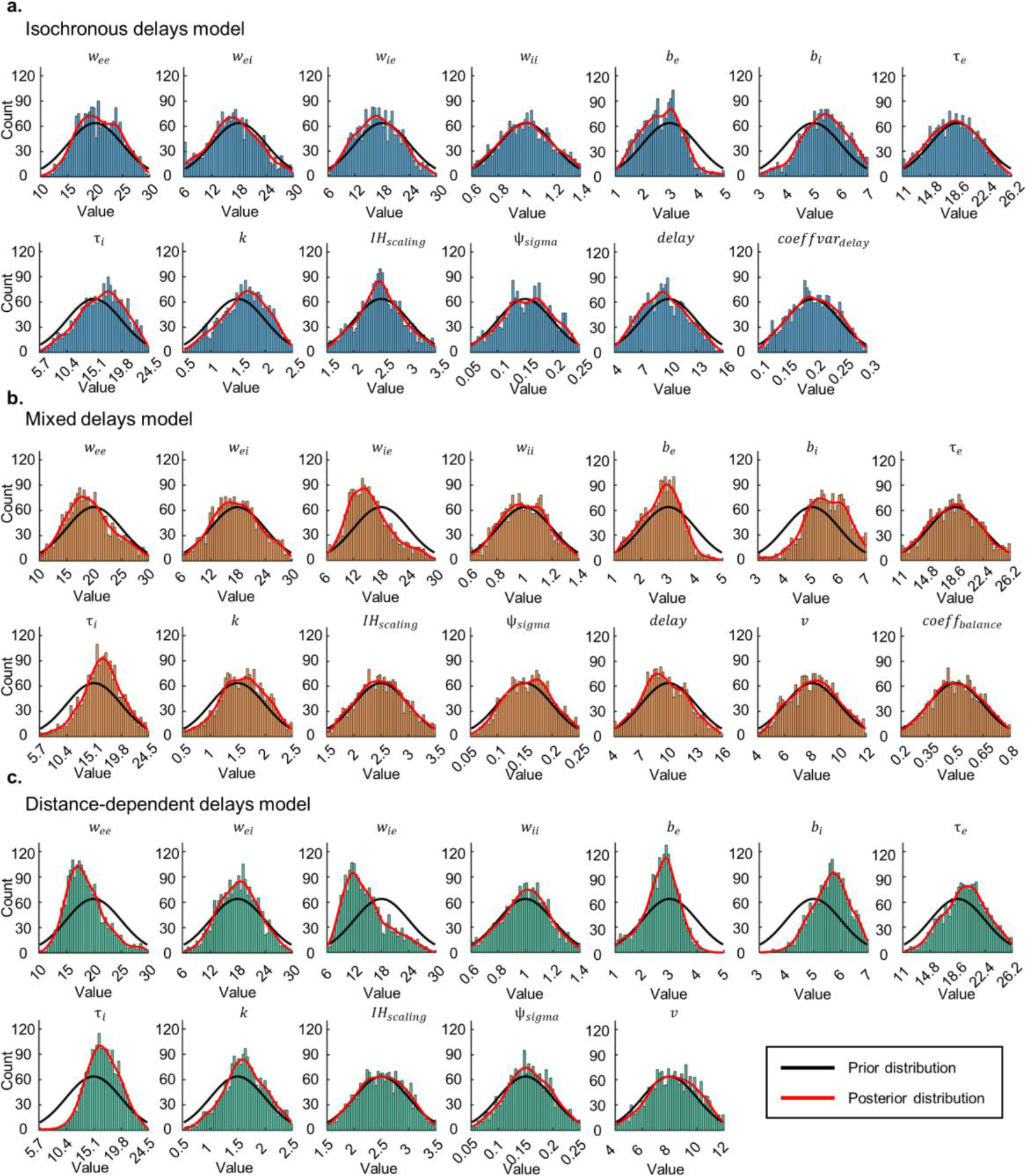
BOLFI yields BNM parameter estimates informed by MEG data a. Marginal posterior distributions of BNM with “isochronous delays” **b.** Marginal posterior distributions of BNM with “mixed delays” **c.** Marginal posterior distributions of BNM with “distance-dependent delays”. Black lines indicate prior distributions, while red lines indicate kernel density estimates of posterior distributions.

We note that the estimated posterior distributions could be used to specify BNMs in future modelling efforts. We refer the reader to our open dataset (Williams et al. (2023)), where we have made available the joint posterior distribution of each of the three BNMs, from which the marginal distributions that we report here, as well as their conditional distributions and joint distributions can be used to specify values of BNM parameters.

### 3.6 Fitted BNM dynamics correspond to those observed in MEG data

The procedure we used to compare the three BNMs assumed the absence of computational problems when the BNMs were fit to MEG data. Hence, we used Posterior Predictive Checks (Gelman et al. (2020), van de Schoot et al. (2021)) to evaluate the fitted BNMs, before comparing them in the next stage. In addition, we used the Posterior Predictive Checks to assess the similarity of the phase synchronization phenomena generated by the three BNMs, when these BNMs were constrained by their respective posterior distributions. We performed the Posterior Predictive Checks by comparing sample medians of four test statistics that we estimated from 1,000 simulations of each of the fitted BNMs, against the value of those test statistics estimated on the MEG dataset. Identical to the Prior Predictive Checks (Section 3.3), the test statistics that we used were the median and median absolute deviation (MAD) of phase synchronization strengths between all region pairs, to measure their central tendency and dispersion respectively. We also estimated the mean and standard deviation of the Kuramoto order parameter, to measure overall strength and variability of zero-lag phase synchronization respectively (see Section 2.3.2 and Table 1 for details of each test statistic). We found the sample medians of the four test statistics estimated on the dynamics of all three BNMs to correspond closely to the values of those test statistics on the MEG dataset (Figure 7a–l). Just as for the Prior Predictive Checks, the value of 0.02 for dispersion in phase synchronization strengths in the MEG dataset was close to the median values of 0.03 for this test statistic across the three methods (Figure 7d–f). In contrast to the Prior Predictive Checks however, the value of 0.09 for central tendency in phase synchronization strengths in the MEG dataset was close to the values of 0.35, 0.27 and 0.17 for this test statistic, for the “isochronous delays”, “mixed delays”, and “distance-dependent delays” methods respectively (Figure 7a–c). The corresponding values from the Prior Predictive Checks were 0.7, 0.78 and 0.79. These results suggest that compared to the mean strengths of phase synchronization generated by the BNMs before fitting, those generated by the fitted BNMs were more similar to those we observed in the MEG dataset while also being more different across BNMs. Just as for the Prior Predictive Checks, the mean and standard deviation of the Kuramoto order parameter had a bimodal distribution for the set of values estimated from the dynamics of all three BNMs. In contrast to the Prior Predictive Checks however, the sample medians of these test statistics were close to their values in the MEG dataset. The value for mean of the Kuramoto order parameter was 0.06, 0.05 and 0.05 for the “isochronous delays”, “mixed delays”, and “distance-dependent delays” methods respectively, which was close to 0.05 for this test statistic in the MEG dataset (Figure 7g–i). Similarly, the value for standard deviation of the Kuramoto order parameter was 0.02 across the three methods, close to the value of 0.03 for this test statistic in the MEG dataset (Figure 7j–l). Compared to the values of the test statistics estimated on the dynamics of the BNMs before fitting, their values estimated on the dynamics of the fitted BNMs corresponded more closely to the values of those test statistics in the MEG dataset. This suggests that compared to the prior distributions of the BNM parameters, their posterior distributions more accurately reflected the ground-truth values of the BNM parameters. Hence, the Posterior Predictive Checks suggested that all three BNMs were fit to the MEG data without computational problems, and that they could be used to choose between the three methods with ABC model comparison.

**Figure 7.**
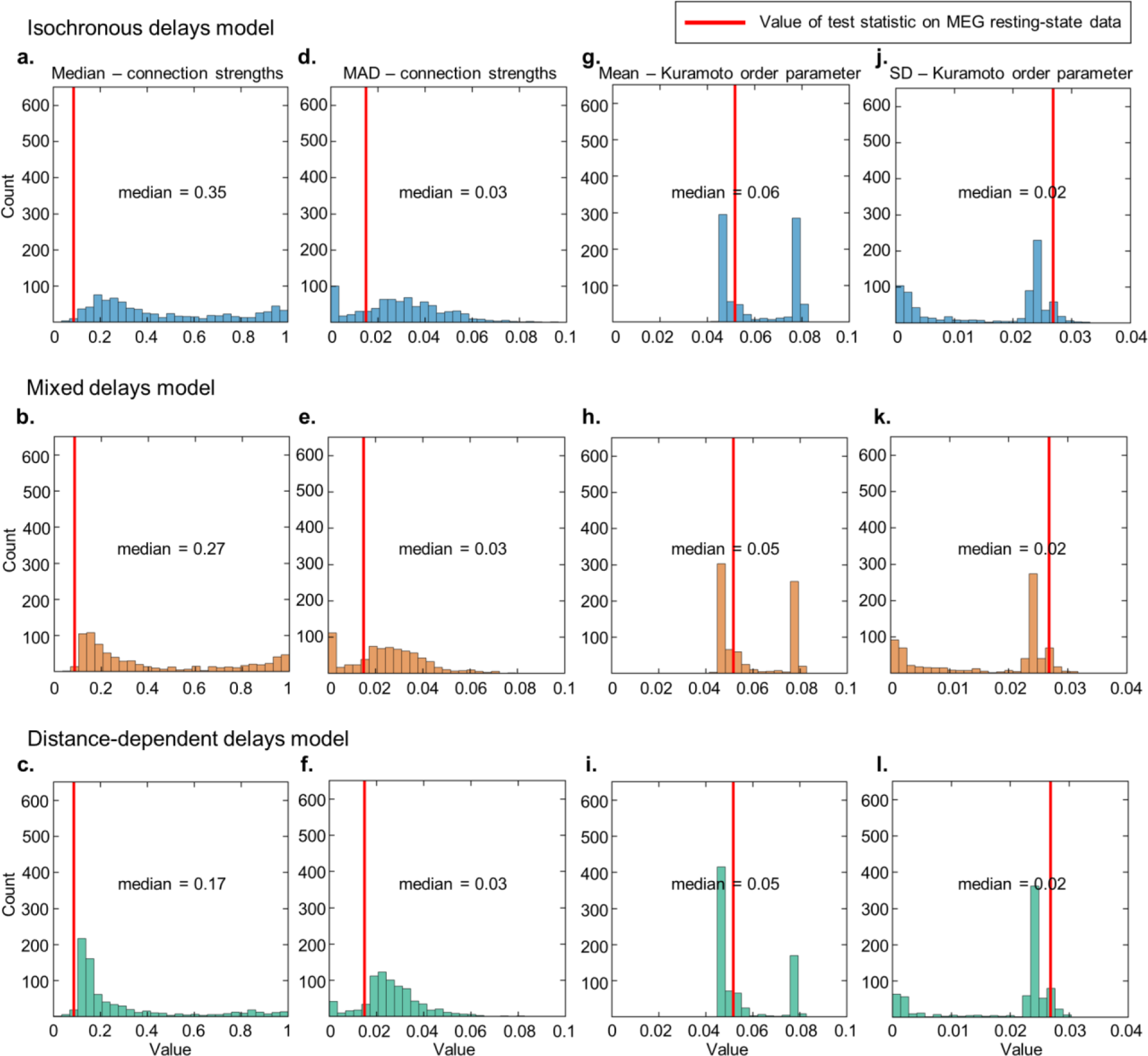
Fitted BNM dynamics correspond to those observed in MEG data. a-c. Histograms of median of alpha-band phase synchronization strengths from multiple BNM simulations, where parameter values were drawn from joint posterior distributions of BNMs with “isochronous delays”, “mixed delays”, and “distance-dependent delays” respectively. **d-f.** Histograms of median absolute deviation (MAD) of alpha-band phase synchronization strengths from multiple BNM simulations, of the three BNMs respectively. **g-i.** Histograms of mean of Kuramoto order parameter from multiple BNM simulations, of the three BNMs respectively. **j-l.** Histograms of standard deviation (SD) of Kuramoto order parameter from multiple BNM simulations, of the three BNMs respectively. For all panels, the red line indicates the corresponding value of that test statistic estimated from the MEG dataset.

### 3.7 BNM with “distance-dependent delays” more probable than BNMs with “isochronous delays” and “mixed delays”

Finally, we compared the three methods to specify inter-regional delays in BNMs of large-scale networks of phase synchronization observed in MEG resting-state. Having fitted BNMs implementing each of the methods to an MEG dataset (*N* = 75), we used ABC model comparison (Beaumont (2019), Sunnåker et al. (2013)) to choose between the fitted BNMs with a separate MEG dataset (*N* = 30). To do so, we first ran 1,000 simulations of each of the three BNMs, with parameter values drawn from their respective joint posterior distributions. For each of the three BNMs, we estimated discrepancies between BNM dynamics from each of the simulations, and MEG data. We computed discrepancy as *ln*(1 − *SSI*)) between vectors of phase synchronization strengths from BNM dynamics and MEG data. We then estimated model probability of each BNM by the relative acceptance rate of discrepancies associated with that BNM, with respect to a range of minimum discrepancies from -1 to 0. Model probabilities represented how likely each of the BNMs were, to describe the generation of large-scale, alpha-band, networks of phase synchronization seen in MEG resting-state data.

The multiple simulations yielded 807, 767 and 779 completed simulations for BNMs with “isochronous delays”, “mixed delays”, and “distance-dependent delays” respectively, the others exceeding the time limit or crossing the memory limit. The model comparison method assumes equal numbers of simulations across BNMs, so we used only the first 767 completed simulations of the three BNMs, *i.e.,* lowest number of completed simulations across BNMs. Model probabilities of the BNM with “distance-dependent delays” were higher than those of the BNMs with “isochronous delays” and “mixed delays”, across thresholds from -0.7 to -0.2 (Figure 8). For a threshold of -0.5 for example, the BNM with “distance-dependent delays” had a probability of 0.54, while the BNM with “mixed delays” had a probability of 0.32 and the BNM with “isochronous delays” had the lowest probability of 0.14. We found the higher probabilities of the BNM with “distance-dependent delays” to be driven by the similarity between its mean strengths of phase synchronization to that observed in the MEG data, rather than the similarity in its standard deviation or its pattern of phase synchronization strengths to those observed empirically (Figure S6). We found the three BNMs to be similarly probable at low thresholds close to 0 and high thresholds close to 1. However, the very low and very high numbers of accepted simulations at these thresholds respectively, render their probabilities non-informative. Notably, we observed an identical pattern of results at intermediate discrepancy thresholds when using eyes-closed MEG resting-state data in the ABC model comparison, inspite of the BNMs being fit to eyes-open MEG resting-state data (Figure S7). Hence, the ABC model comparison revealed the BNM with “distance-dependent delays” as the most probable and the BNM with “isochronous delays” as the least probable, of describing the generating of large-scale networks of phase synchronization seen in MEG.

**Figure 8.**
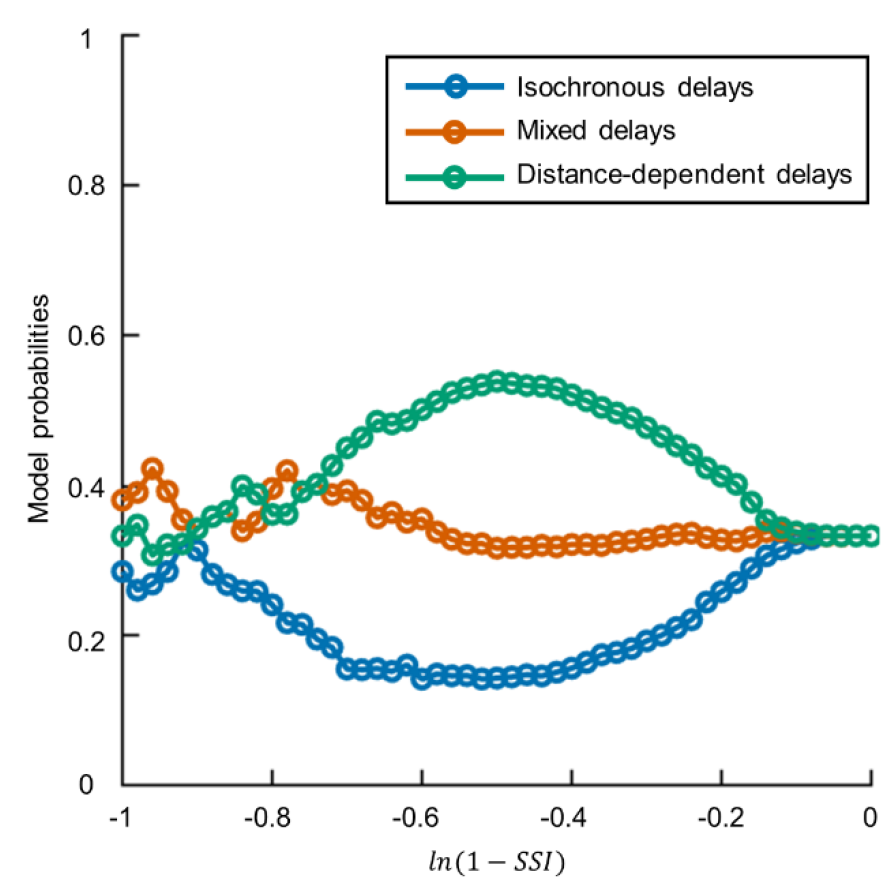
BNM with “distance-dependent delays” more probable than BNMs with “isochronous delays” and “mixed delays”. Model probabilities of BNMs with “isochronous delays”, “mixed delays”, and “distance-dependent delays”, for a range of minimum discrepancies between phase synchronization strengths of BNM dynamics and MEG data. Discrepancies are estimated as *ln*(1 − *SSI*)), where SSI is the Structural Similarity Index.

While the three BNMs differed in the extent to which distance between brain regions determined the inter-regional delays, they also differed in the variability or heterogeneity of their delays. BNMs with “distance-dependent delays” had the highest delay heterogeneity. Hence, we performed a control analysis to assess whether the correspondence between phase synchronization strengths of the BNM with “distance-dependent delays” and those in MEG data arose merely from delay heterogeneity. To do so, we ran 1,000 simulations of a BNM with “randomised delays”, where we used randomly resampled (without replacement) versions of “distance-dependent delays” used in the ABC model comparison. We simulated the BNM with “randomised delays” at the same parameter values we had used to simulate the BNM with “distance-dependent delays” in the ABC model comparison. Then, we estimated discrepancies between dynamics of the BNMs with “randomised delays” and MEG data, and used a Wilcoxon rank-sum test to compare these to the corresponding discrepancies for the BNMs with “distance-dependent delays”. We found the discrepancies for the BNM with “distance-dependent delays” to be much lower (*p* = 4.5*e*^-46^) than those for the BNM with “randomised delays” (Figure S8). The sample median of discrepancies for the BNM with “distance-dependent delays” was -0.41, while the sample median of discrepancies for the BNM with “randomised delays” was -0.22. Hence, the control analysis revealed that mere delay heterogeneity does not account for the correspondence between inter-regional phase synchronization strengths of the BNM with “distance-dependent delays” and those in MEG data. These results rule out alternative explanations for the BNM with “distance-dependent delays” being more probable than BNMs with “isochronous delays” and “mixed delays”, of describing the generation of large-scale networks of phase synchronization seen in MEG.

## 4. Discussion

Large-scale networks of phase synchronization are considered to regulate communication between brain regions, but the relationship to their structural substrates remains poorly understood. In this study, we used an ABC workflow to compare the “isochronous delays”, “mixed delays”, and “distance-dependent delays” methods of specifying inter-regional delays in BNMs of phase synchronization. Prior Predictive Checks revealed BNMs of all three methods to generate phase synchronization phenomena encompassing those observed in MEG resting-state. Fitting the BNMs to MEG resting-state data yielded reliable posterior distributions of parameters of all the three BNMs. Finally, ABC model comparison of the fitted BNMs revealed the BNM with “distance-dependent delays” to be the most probable to describe the generation of large-scale networks of phase synchronization seen in MEG.

Previous modelling studies have demonstrated the role of distance-dependent inter-regional delays in generating power spectra of MEG activity from individual brain regions (Cabral et al. (2022)), alpha-band inter-regional networks of amplitude correlation (Cabral et al. (2014), Nakagawa et al. (2014)), and the observed bimodal distribution (Dotson et al. (2014), Dotson et al. (2015)) in angles of inter-regional phase synchronization (Petkoski et al. (2018), Petkoski & Jirsa (2019)). However, networks of phase synchronization are physiologically distinct from networks of amplitude correlation (Engel et al. (2013)) and exhibit different patterns of connectivity (Siems & Siegel (2020)). Similarly, angles of phase synchronization are distinct from the strengths of phase synchronization that we modelled. In contrast to these studies, we demonstrated the role of distance-dependent conduction delays in generating alpha-band inter-regional networks of phase synchronization observed in MEG resting-state.

Those few modelling studies which use BNMs with distance-dependent delays to generate networks of phase synchronization only contrast them to BNMs with “zero delays”. These studies (Abeysuriya et al. (2018), Finger & Bönstrup et al. (2016)) have demonstrated BNMs with distance-dependent delays to generate alpha-band networks of phase synchronization more similar to those in MEG or EEG resting-state, than networks from BNMs with “zero delays”. However, “zero delays” are biologically implausible, implying infinite conduction velocities. In contrast, we demonstrate that BNMs with distance-dependent delays generate networks more similar to those in MEG resting-state than those from BNMs implementing two biologically plausible methods accounting for spatially varying conduction velocities.

The generation of phenomena observed in MEG, *e.g.*, power spectra, amplitude correlations (Cabral et al. (2022), Cabral et al. (2014)) by BNMs with distance-dependent delays has been linked to the variability or heterogeneity of these delays (Lee et al. (2009), Touboul (2012)). We demonstrate that inter-regional distances rather than delay heterogeneity *per se*, explain the similarity between alpha-band networks of phase synchronization generated by BNMs with distance-dependent delays, and those observed in MEG resting-state.

Previous neurophysiological and modelling studies have contributed to our understanding of the structure-function relationship underlying phase synchronization. For example, studies have demonstrated the role of excitatory-inhibitory connections in generating local oscillatory dynamics (Buzsáki (2006), Traub et al. (1997)) required for phase synchronization, and the role of anatomical connections in promoting inter-regional phase synchronization (Gray (1994), Finger & Bönstrup et al. (2016)). In our study, intermediate diagnostics from BOLFI model fitting corroborated these previous results. For example, we found supporting evidence for the role of intra-regional connections between excitatory and inhibitory populations in generating local oscillatory dynamics, and for the role of inter-regional anatomical connections in promoting inter-regional phase synchronization. In addition to these previous studies, we furnish new understanding on the role of inter-regional delays in generating large-scale networks of phase synchronization observed in MEG resting-state. Our results suggest that the dynamics of brain regions interact though inter-regional anatomical connection via distance-dependent delays to generate large-scale networks of phase synchronization.

Inter-regional conduction delays reported in human and animal neurophysiological studies provide a basis for comparison to the distance-dependent conduction delays suggested by our modelling study. Human studies have reported correlations of 0.44 between tract length and the onset latency of the stimulation-based evoked potential in intra-cranial EEG recordings (Trebaul et al. (2018)), which is consistent with the linear relationship between inter-regional distance and inter-regional delays suggested by our study. Inter-regional delays estimated with a model-based approach on intra-cranial EEG recordings (Lemaréchal et al. (2022)) also reported a linear relationship between tract length and estimated delays for most brain regions, consistent with the distance-dependent delays suggested by our study.

In contrast to the distance-dependent conduction delays reported for most brain regions with intra-cranial EEG recordings (Lemaréchal et al. (2022)), some brain regions present highly similar conduction delays with several other regions. For example, the right insula has highly similar conduction delays between 6–8 ms with several ipsilateral brain regions (Lemaréchal et al. (2022)). Animal neurophysiological studies have also presented evidence for isochronous delays, in specific brain regions. For example, efferent connections of layer V neurons from regions in the rat ventral temporal cortex had largely isochronous conduction delays with several ipsilateral brain regions (Chomiak et al. (2008)), and afferent connections of layer IV neurons from thalamus also had highly similar delays with a number of cortical brain regions (Salami et al. (2003)). These highly similar delays for a few brain regions might be due to regulation in conduction velocities by activity-dependent myelination (Noori et al. (2020)), in response to specialised roles of these regions in functions involving fine temporal coordination, *e.g.*, sensory cue processing (Chomiak et al. (2008), Pajevic et al. (2014)). We propose that future work could investigate methods to specify inter-regional delays, which account for the region-specific nature of their distance-dependence.

Due to their high delay heterogeneity, BNMs with distance-dependent delays might be prone to the dynamical regime of amplitude death, *i.e.*, cessation of oscillations (Atay (2003)).

Phase synchronization cannot occur in regimes of amplitude death due to absence of oscillations and in fact, dynamically adjusting conduction velocities by activity-dependent myelination regulation has been suggested as a means of avoiding this regime (Pajevic et al. (2014)). However, our Posterior Predictive Checks revealed oscillatory dynamics from several simulations of the fitted BNM with distance-dependent delays, despite the highly heterogeneous nature of these delays. Further, BNMs from previous studies (Cabral et al. (2022)) report regimes of reduced amplitude rather than amplitude death, despite using distance-dependent conduction delays which are highly heterogeneous by nature.

Distance-dependent conduction delays predict long inter-regional delays between spatially distant brain regions. These long delays might be deleterious to inter-regional communication through phase synchronization, particularly when the lag of phase synchronization is close to the oscillatory time period (Aboitiz et al. (2003), Pajevic et al. (2014)). However, modelling studies have demonstrated several means by which phase synchronization lags might be adjusted, enabling rapid inter-regional communication despite long conduction delays. For example, the presence of a common relay region between two interacting regions (Vicente et al. (2008)), driving currents (Tiesinga et al. (2010)), or local inhibition (Battaglia et al. (2007)) can adjust the lag of phase synchronization towards zero. Hence, temporally precise inter-regional communication can occur despite the presence of long inter-regional delays.

We mention some limitations of our study and propose approaches to addressing these. First, we used the Euclidean distance between regions divided by conduction velocity to estimate inter-regional delays. Using Euclidean distance to specify tract length facilitated comparison to several previous modelling studies on brain functional networks (Abeysuriya et al. (2018), Hadida et al. (2018), Cabral et al. (2014), Nakagawa et al. (2014), Deco et al. (2009), Ghosh et al. (2008)), which also used this measure. However, any spatially varying errors in tract length estimation introduced by Euclidean distance could mask the contribution of spatially varying conduction velocities in determining inter-regional delays. Diffusion MRI-based tractography can potentially provide more accurate estimates of the tract length, but current methods are also prone to error from seeding and termination biases (Girard et al. (2014), Sotiropoulos & Zalesky (2019)). Future work could employ the ABC workflow we used, to compare different methods to specify tract lengths, thereby further constraining BNMs of inter-regional networks of phase synchronization. Second, we focused only on alpha-band frequencies due to the clear evidence for alpha-band oscillations both in our own MEG dataset and in previous MEG resting-state studies (Mahjoory et al. (2020)), oscillations being a pre-requisite for phase synchronization. Hence, our findings are only relevant to phase synchronization in alpha-band frequencies. However, we note that brain regions also generate oscillatory activity in delta, low-beta and high-beta frequency bands (Mahjoory et al. (2020)). Future modelling work could study phase synchronization in multiple frequency bands by, *e.g.*, including multiple generators per brain region (Deco et al. (2017)). Note that broadening the range of frequencies studied would change the values of the summary statistics we use to describe the BNM dynamics and those in MEG data, likely resulting in changes to the posterior distributions of the BNM parameters to those we have reported here. Third, we assumed that all brain regions generate oscillations, in line with empirically observed cortex-wide alpha-band spectral peaks both in our own MEG dataset and in previous MEG resting-state studies (Mahjoory et al. (2020)). However, we acknowledge recent evidence from intra- cranial EEG and MEG data suggesting that not all brain regions might generate oscillations (Myrov et al. (2023)). Future modelling studies could examine the role of sparse oscillation generators across cortex, including the interaction between sparsity and inter-regional delays, in the structure-function relationship of large-scale networks of phase synchronization.

Finally, we assumed BNM parameters governing local dynamics to be identical across brain regions. This was effective in limiting the number of BNM parameters to be estimated, while introducing region-wise variation in BNM parameters would have exponentially increased the volume of parameter space resulting in much higher numbers of BNM simulations required to sample the parameters space (Gutmann & Corander (2016)). However, we acknowledge empirical evidence for region-wise variation in structural and functional properties of brain regions (Markello & Hansen et al. (2022)), and modelling work suggesting the utility of informing BNMs with this region-wise variation (Demirtaş et al. (2019), Sanz-Perl et al. (2022)) to emulate empirically observed dynamics. Future BNMs of inter-regional phase synchronization could parameterise region-wise variation with only a few parameters, by, *e.g.*, expressing the variation in terms of empirically observed spatial gradients (Mahjoory et al. (2020), Markello & Hansen (2022)).

The ABC workflow that we employed for model fitting and model comparison naturally accounted for uncertainty in values of BNM parameters. We also employed a number of recommended best practices (Gelman et al. (2020), van de Schoot et al. (2021)) as we proceeded from specifying the three BNMs through to fitting these BNMs and comparing the fitted BNMs. In particular, we i) used prior distributions of BNM parameters informed by the aggregated neurophysiology literature and the literature on modelling brain functional networks, ii) verified suitability of the prior distributions and specification of the BNMs with Prior Predictive Checks, iii) verified assumptions underlying the BOLFI model fitting with fake-data simulations, iv) used diagnostics of the GP-based surrogate modelling to assess intermediate stages of the BOLFI model fitting, v) employed established convergence diagnostics to assess reliability of the estimated posterior distributions of BNM parameters, vi) verified that BOLFI fitting had completed without error using Posterior Predictive Checks, vii) performed ABC model comparison across a range of discrepancy thresholds, and completed a control analysis to rule out alternative explanations for the results of the ABC model comparison. We therefore propose that our results are robust. In conclusion, we found evidence that distance-dependent delays crucially contribute to the generation of alpha-band inter-regional networks of phase synchronization observed in MEG resting-state.

## Acknowledgements

The authors are grateful to the reviewers of this manuscript for their thoughtful comments, addressing which has substantively improved the manuscript. Further, we acknowledge Finnish Centre for Artificial Intelligence (FCAI), Academy of Finland (NW: 321542, SK: 292334, 956 319264, MP: 253130, 256472, 281414, 296304, 266745, SP: 266402, 266745, 303933, 957 325404), Department of Science & Technology (DST), India and Sigrid Juselius Foundation, for providing funding for this project. The authors are grateful to Prof. Sitabhra Sinha, Dr. Chandrasekhar Kuyyamudi, Dr. Ayush Bharti, Dr. Henri Pesonen, Dr. Michael Gutmann and Antti Karvanen for invaluable discussions, and to Alex Aushev, Anirudh Jain, Diego Mesquita and Sophie Wharrie for comments on manuscript drafts. Most of all, we are very grateful to Jarno Rantaharju, Thomas Pfau, Richard Darst and Enrico Glerean from Aalto Scientific Computing, for facilitating use of computational resources provided by the Aalto Science-IT project.

## Supplementary figures

**Figure S1.**
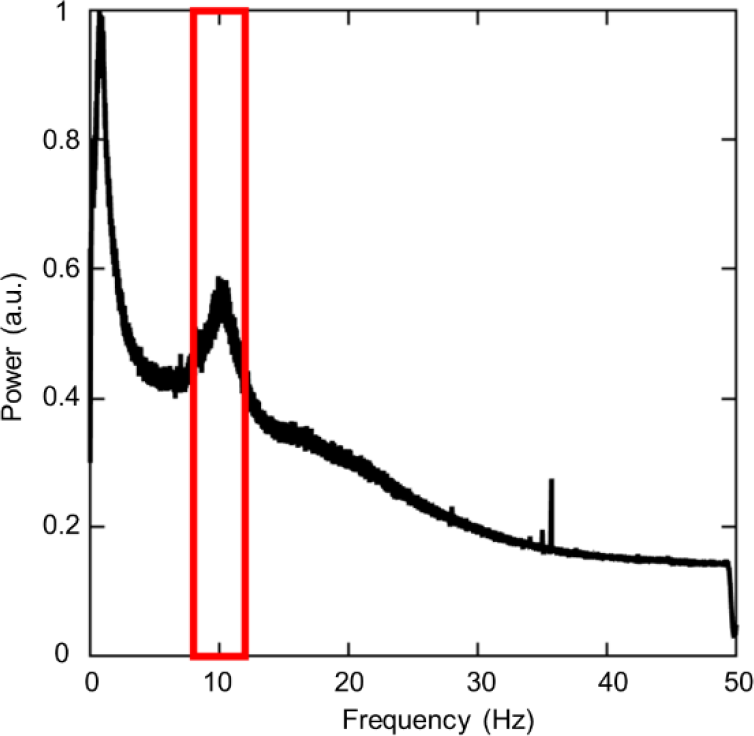
Group-level frequency spectrum displays alpha-band spectral peak. Group-level frequency spectrum averaged across brain regions, of source-reconstructed MEG resting-state data. Red borders of rectangle outline band of alpha frequencies (8–12 Hz).

**Figure S2.**
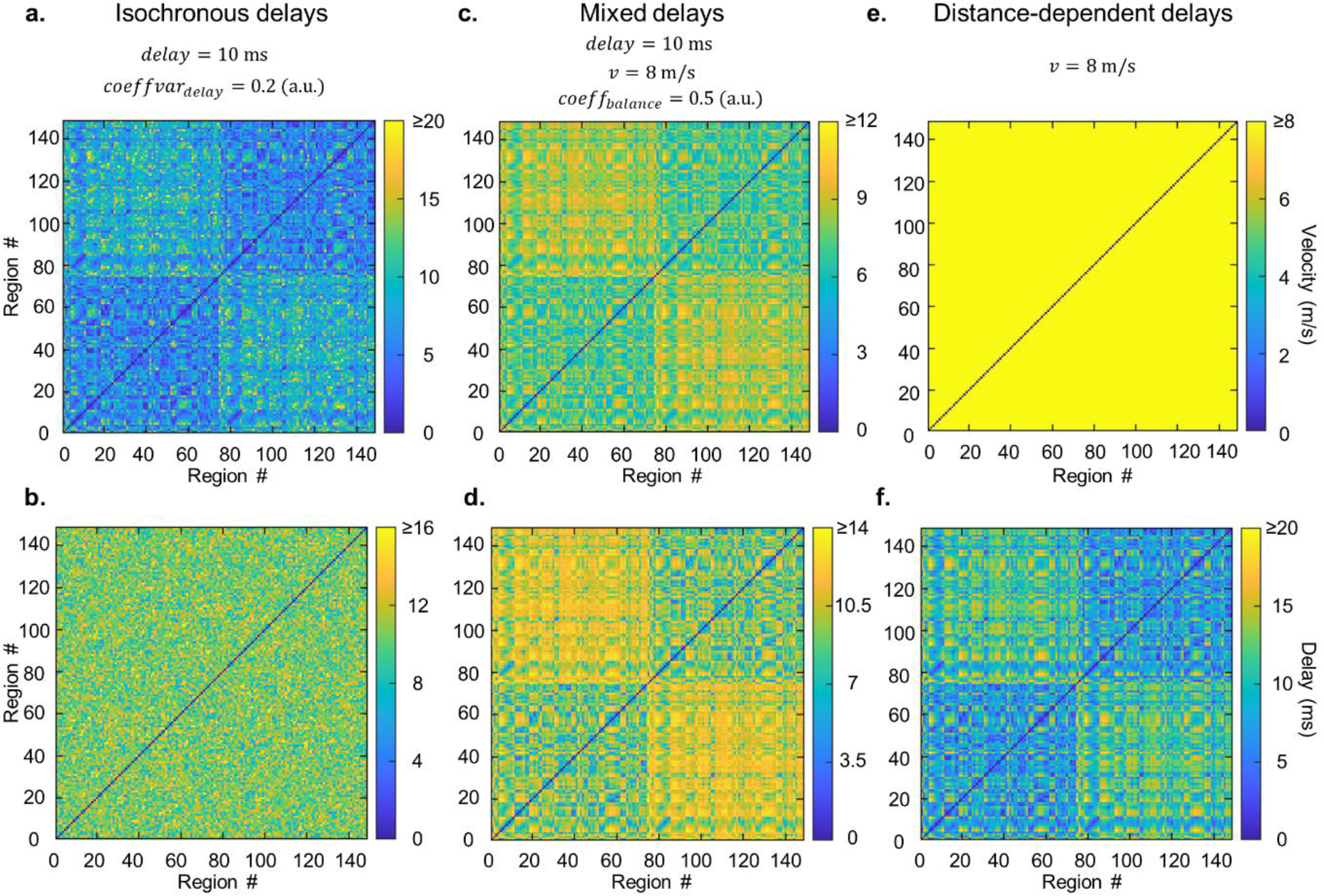
Specifying inter-regional delays for “isochronous delays”, “mixed delays” and “distance-dependent delays” methods. a-b. Matrices of inter-regional conduction velocities and conduction delays for “isochronous delays” method, **c-d.** Matrices of inter-regional conduction velocities and conduction delays for “mixed delays” method, **e-f.** Matrices of inter-regional conduction velocities and conduction delays for “distance-dependent delays” method.

**Figure S3.**
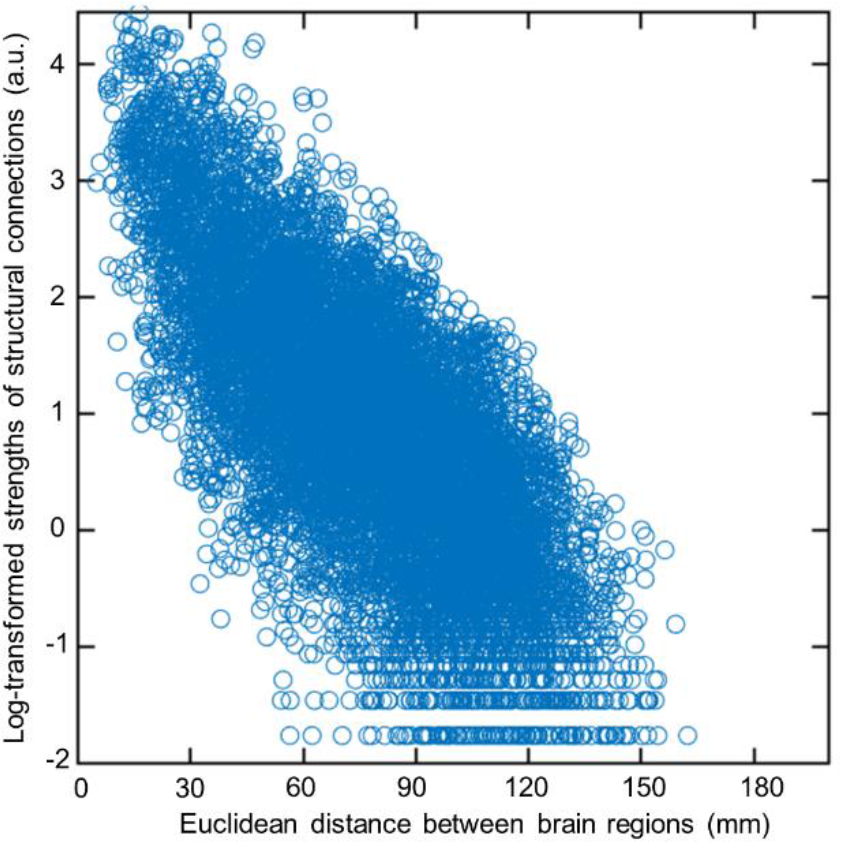
Log-transformed strengths of structural connections are inversely related to distance between brain regions. Scatter plot of Euclidean distance between every pair of brain regions in the 148-region Destrieux brain atlas and log-transformed strengths of structural connections between these regions.

**Figure S4.**
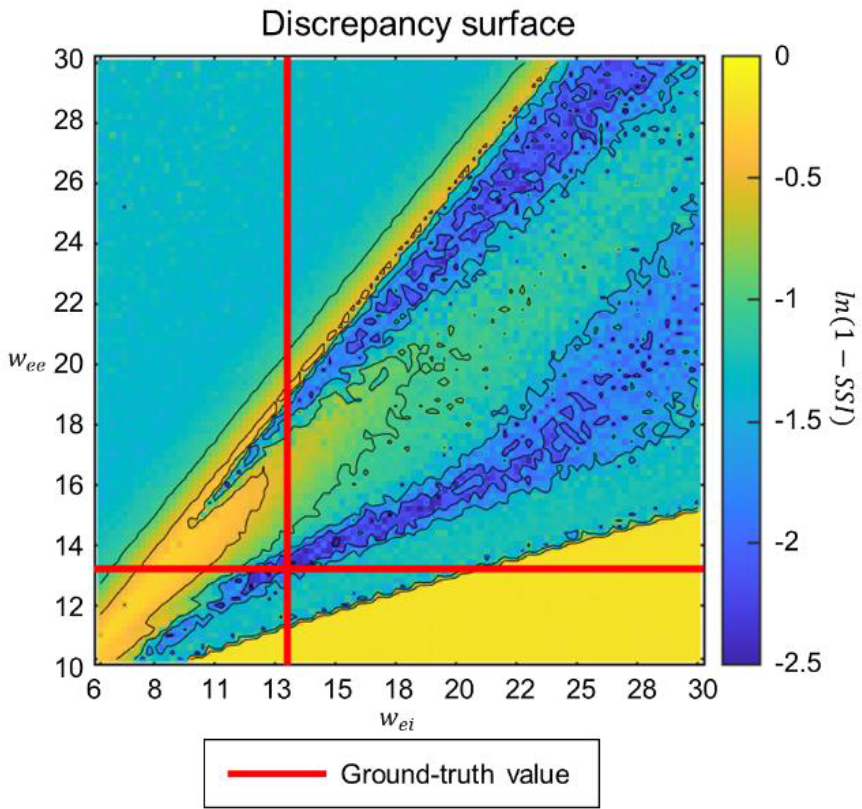
Gaussian Process (GP)-predicted discrepancies are sensitive to values of BNM parameters. 100 ×100 grid of GP-predicted discrepancies between summary statistics of BNM dynamics at every pair of *w*_*ee*_ and *w*_*ei*_ values, and BNM dynamics at *w*_*ee*_ =12.9, *w*_*ei*_=13.4. *w*_*ee*_ is the strength of connections within excitatory neuronal populations, *w*_*ei*_ is the strength of connections from inhibitory to excitatory neuronal populations. Red lines indicate ground-truth values of *w*_*ee*_ and *w*_*ei*_ . Discrepancies were measured by *ln*(1 − *SSI*)). SSI is the Structural Similarity Index.

**Figure S5.**
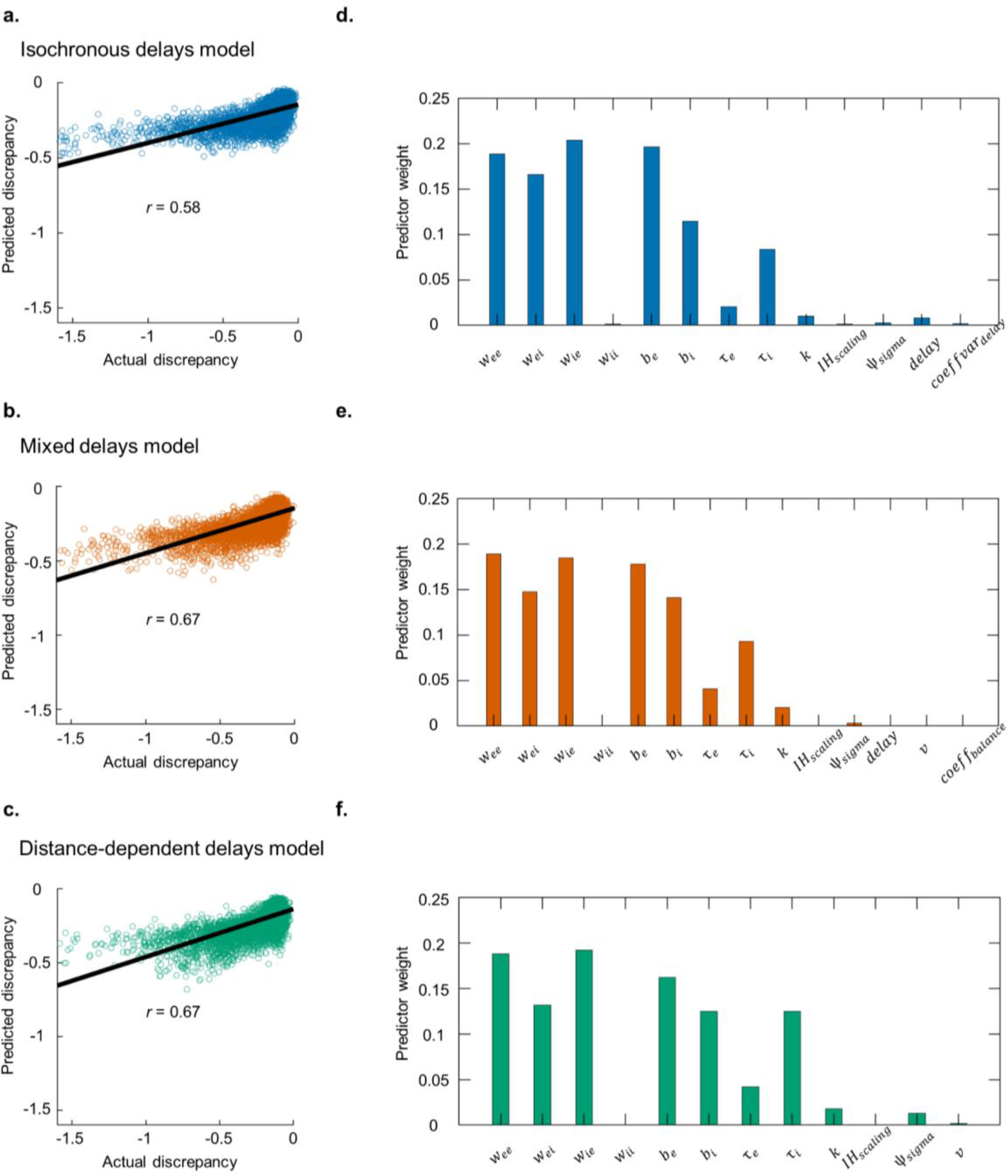
Gaussian Process (GP) regression results during application of BOLFI to fit BNMs to MEG resting-state data. a-c. Scatter plot of actual and GP-predicted discrepancies between BNM dynamics and MEG data, for the “isochronous delays”, “mixed delays” and “distance-dependent delays” methods respectively. **d-f** Relative importance of each BNM parameter in predicting discrepancies BNM dynamics and MEG data, for the three methods.

**Figure S6.**
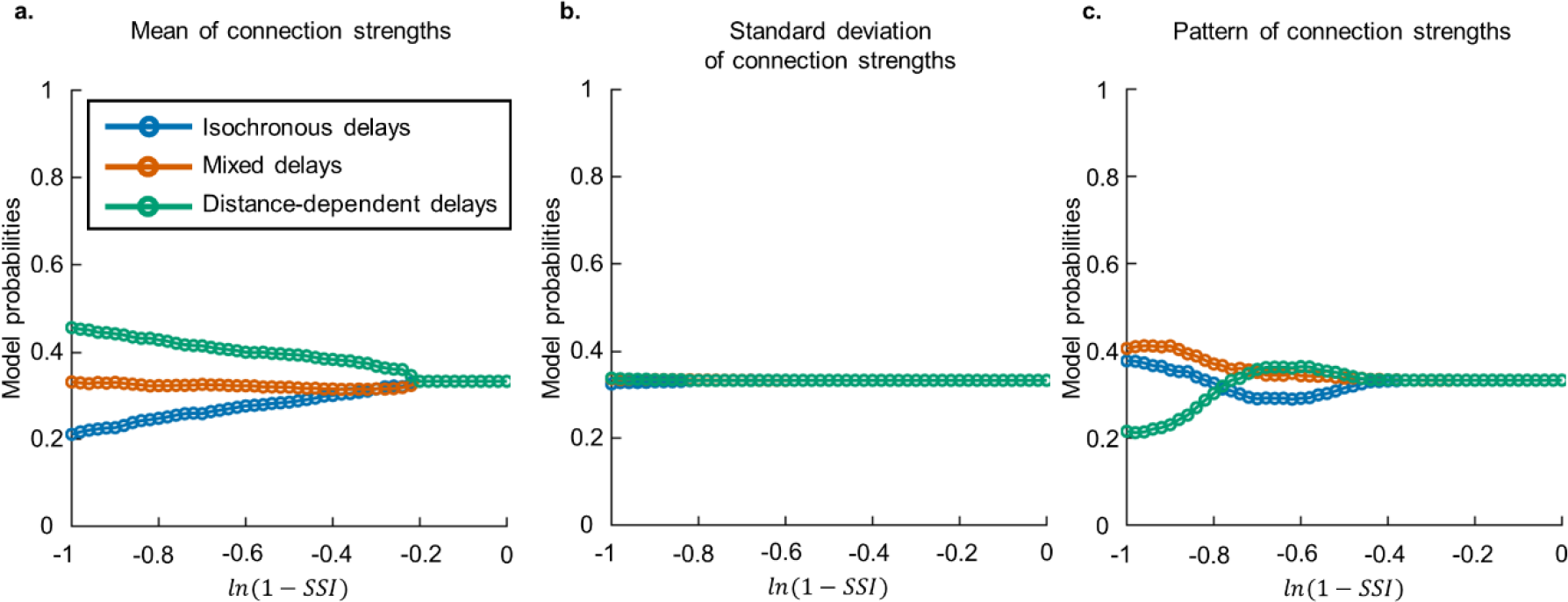
BNM with “distance-dependent delays” most probable when comparing mean of connection strengths, but not their standard deviation or pattern. a. Model probabilities of BNMs with “isochronous delays”, “mixed delays” and “distance-dependent delays”, when comparing mean of connection strengths. **b.** Model probabilities of BNMs with “isochronous delays”, “mixed delays” and “distance-dependent delays”, when comparing standard deviation of connection strengths. **c.** Model probabilities of BNMs with “isochronous delays”, “mixed delays” and “distance-dependent delays”, when comparing pattern of connection strengths. Discrepancies are estimated as *ln*(1 − *SSI*)), where SSI is the Structural Similarity Index.

**Figure S7.**
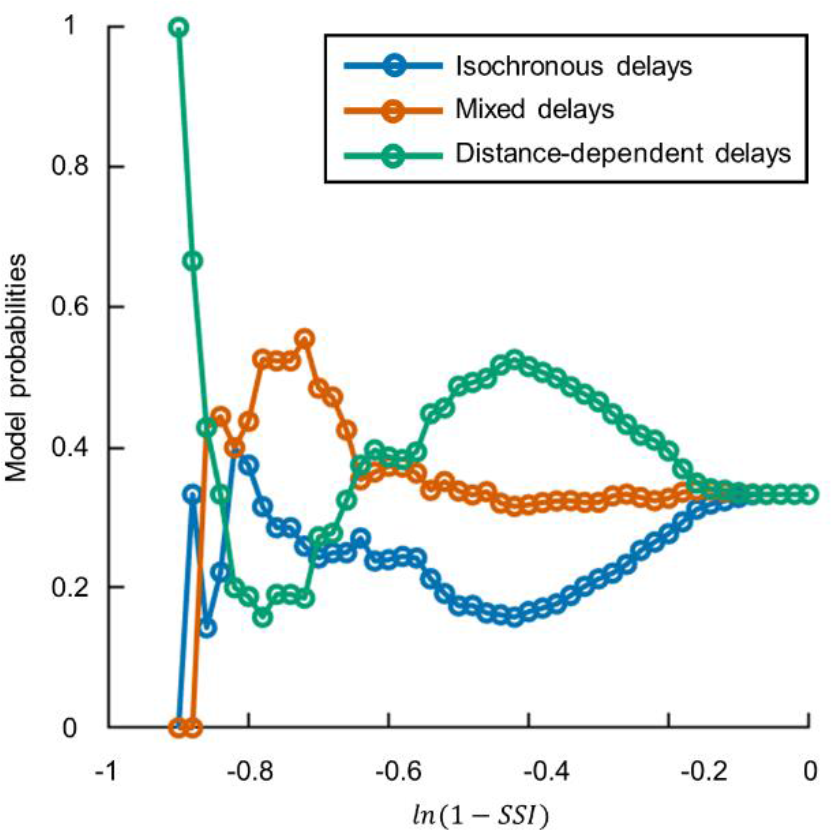
BNM with “distance-dependent delays” more probable than BNMs with “isochronous delays” and “mixed delays” for eyes-closed MEG resting-state data. Model probabilities of BNMs with “isochronous delays”, “mixed delays”, and “distance-dependent delays”, for a range of minimum discrepancies between phase synchronization strengths of BNM dynamics and MEG eyes-closed resting-state data. Discrepancies are estimated as *ln*(1 − *SSI*)), where SSI is the Structural Similarity Index.

**Figure S9.**
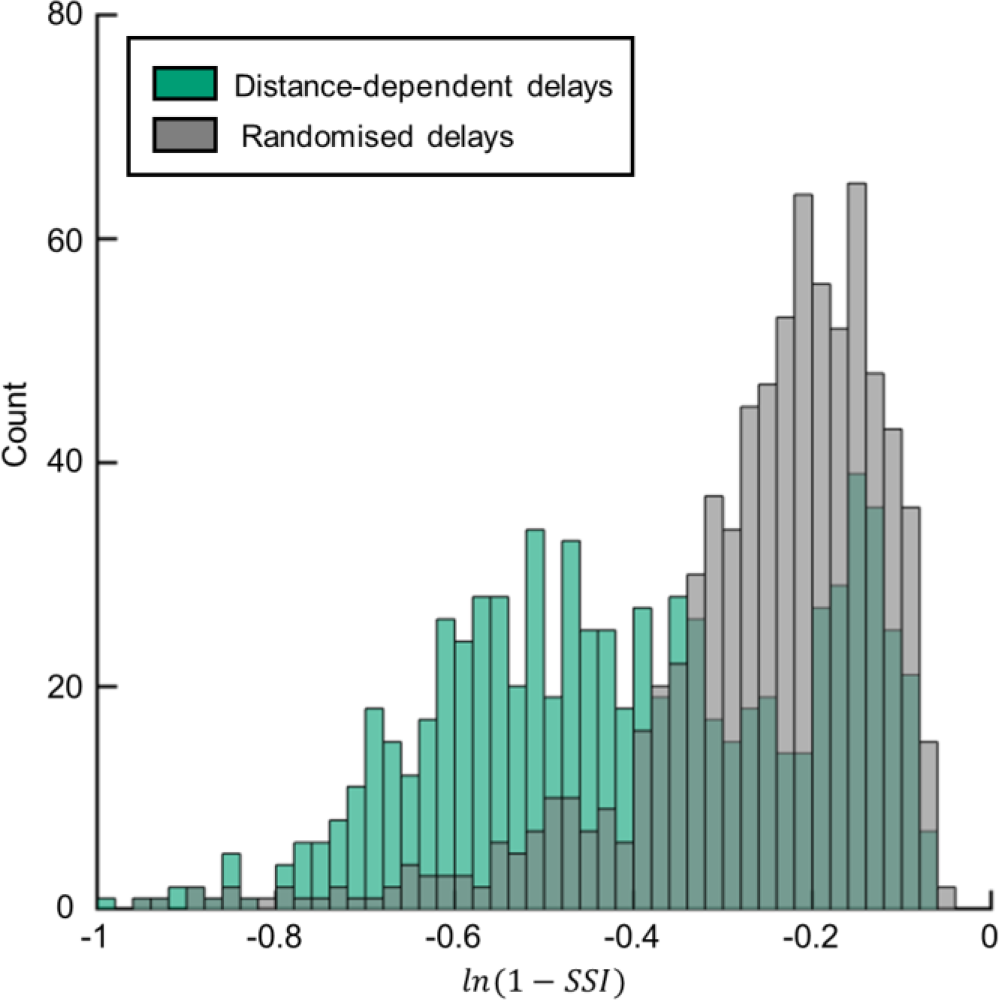
Delay heterogeneity does not explain correspondence between phase synchronization strengths of dynamics from BNM with “distance-dependent delays” and those in MEG data. Histogram of discrepancies between dynamics of BNM with “distance-dependent delays” (green) and MEG data, and histogram of discrepancies between dynamics of BNM with “randomised delays” (gray) and MEG data. Histogram overlap is also shown (dark green). Discrepancies are *ln*(1 − *SSI*)), SSI is the Structural Similarity Index.

